# Characterization of ribosome stalling and no-go mRNA decay stimulated by the Fragile X protein, FMRP

**DOI:** 10.1101/2024.02.02.577121

**Authors:** MaKenzie R. Scarpitti, Benjamin Pastore, Wen Tang, Michael G. Kearse

**Affiliations:** Department of Biological Chemistry and Pharmacology, Center for RNA Biology, The Ohio State University, Columbus, OH 43210 USA

**Keywords:** neurological disease, ribosome, RNA binding protein, translation control, translation regulation

## Abstract

Loss of functional fragile X mental retardation protein (FMRP) causes fragile X syndrome (FXS) and is the leading monogenic cause of autism spectrum disorders and intellectual disability. FMRP is most notably a translational repressor and is thought to inhibit translation elongation by stalling ribosomes as FMRP-bound polyribosomes from brain tissue are resistant to puromycin and nuclease treatment. Here, we present data showing that the C-terminal non-canonical RNA-binding domain of FMRP is essential and sufficient to induce puromycin-resistant mRNA•ribosome complexes. Given that stalled ribosomes can stimulate ribosome collisions and no-go mRNA decay (NGD), we tested the ability of FMRP to drive NGD of its target transcripts in neuroblastoma cells. Indeed, FMRP and ribosomal proteins, but not PABPC1, were enriched in isolated nuclease-resistant disomes compared to controls. Using siRNA knockdown and RNA-seq, we identified 16 putative FMRP-mediated NGD substrates, many of which encode proteins involved in neuronal development and function. Increased mRNA stability of the putative substrates was also observed when either FMRP was depleted or NGD was prevented via RNAi. Taken together, these data support that FMRP stalls ribosomes and can stimulate NGD of a select set of transcripts in cells, revealing an unappreciated role of FMRP that would be misregulated in FXS.

## INTRODUCTION

Loss of functional fragile X mental retardation protein (FMRP) causes fragile X syndrome (FXS) (1–4) and is the leading monogenic cause of autism spectrum disorders and intellectual disability (5). FMRP is an RNA-binding protein (RBP) that is highly expressed in the brain and gonads of both males and females (6); however, males are more phenotypically affected due to the X-linked mutation. While the complete molecular mechanism of FMRP has been disputed, most data support that FMRP acts as a translational repressor by inhibiting elongating ribosomes (7–11), yet FMRP may also inhibit initiation of specific mRNAs or in certain contexts (12–14). Thus, at least one facet of the FXS phenotype is believed to be caused by aberrant and hyperactive protein synthesis at neuronal synapses (15–17).

The ability of FMRP to act as an RBP is centered in all proposed models of FMRP function (6). To bind RNA, FMRP harbors at least three RNA-binding elements including two canonical and structured KH1 and KH2 domains, as well as a short RGG box motif within the long flexible C-terminal domain (CTD) (18–20). A third KH domain (i.e., KH0) has been reported based off of structural homology (20) but does not have detectable RNA-binding ability *in vitro* (21). We and others have previously biochemically dissected human FMRP and reported that the RGG box motif and the adjacent CTD together are essential and sufficient to inhibit translation of multiple reporter mRNAs in rabbit reticulocyte lysate (RRL) (8, 9). Importantly, our recent work has shown that the RGG box motif and CTD together, but not separately, form a non-canonical RNA-binding domain (ncRBD) that allows broader RNA-binding ability than previously reported (9). This observation helps explain the inability to identify a consensus motif for FMRP target transcripts in cells.

Consistent with the role of FMRP inhibiting translation elongation in neurons, FMRP is in association with puromycin-and RNase-resistant polyribosome complexes isolate from mouse brain (7). Since the ability of puromycin to act as a polypeptide chain terminator is specific for actively elongating ribosomes or those with an unoccupied A site (22), the puromycin-resistant complexes are thought to be translationally stalled. This is supported by the tight ribosome packing that is resistant to nuclease treatment. In alignment with this model, our recent work has shown that human FMRP inhibits translation of nanoLuciferase (nLuc) reporter mRNA and causes the accumulation of ribosomes on reporter mRNA *in vitro*. Importantly, our *in vitro* FMRP-mediated translational control system faithfully recapitulates the formation of puromycin-resistant ribosomes on reporter mRNAs with human FMRP (9, 23) similar to what has been reported in brain tissue (7).

It is now widely understood that stalled ribosomes often trigger ribosome collisions that subsequently stimulate the no-go mRNA decay (NGD) pathway. In such examples, the leading ribosome is inactive and sterically hinders trailing ribosomes (24, 25). The interface between collided ribosomes forms a unique substrate for the ubiquitin E3 ligase ZNF598 (26) (Hel2 in yeast (27, 28)). After ubiquitylation of small ribosomal subunit proteins RPS10 and RPS20 by ZNF598 (29), an endonuclease (Cue2 in yeast (30), NONU-1 in *Caenorhabditis elegans* (31), and likely N4BP2 in humans (30)), is recruited to cleave the mRNA between collided ribosomes. Ribosome collisions and NGD are generally thought to be a quality control mechanism for cells to identify and destroy damaged or truncated mRNAs that, if translated, may be toxic. However, given that multiple lines of evidence support the ability of FMRP to stall elongating ribosomes, it is plausible, yet untested, that one role of FMRP is to induce ribosome collisions and NGD to negatively regulate gene expression of its target mRNAs.

In this report we test and characterize the ability of FMRP to form puromycin-resistant mRNA•ribosome complexes and to cause ribosome collisions *in vitro* in RRL, as well as to stimulate NGD in neuroblastoma cells. Using a series of biochemical approaches, RNAi, and RNA-seq, the data presented here support that FMRP does stall ribosomes and can stimulate NGD in neuroblastoma cells, albeit on a small number of transcripts, revealing an unappreciated role of FMRP that would be misregulated in FXS when FMRP is lost.

## RESULTS

### The ncRBD of FMRP, composed of the RGG box motif and the CTD, is essential and sufficient to stall ribosomes *in vitro*

Recombinant N-terminally truncated human FMRP (NT-hFMRP) (**Figure 1A, B**) has been shown to be more stable than recombinant full-length FMRP and retains translational repression activity (32). Using NT-hFMRP, we previously reported that FMRP inhibits translation, causes the accumulation of ribosomes on translationally repressed transcripts, and forms puromycin-resistant mRNA•ribosome complexes *in vitro* (9). Additionally, we have detailed that the RGG box motif and the CTD together form a ncRBD which elicits broad RNA-binding ability. Here, we use our published puromycin-induced mRNA•ribosome dissociation assay (9, 23) with nLuc reporter mRNPs formed with WT and mutant FMRP to test and confirm that the translationally repressive ncRBD of FMRP also drives the formation of puromycin-resistant mRNA•ribosome complexes (**Figure 1A, B and Figure S1)**. This assay takes advantage of the tyrosyl-tRNA mimic, puromycin, to act as a chain terminator of translation and to cause dissociation of the mRNA from ribosomes that are actively elongating or are in the unrotated/classic state with an unoccupied A site (22, 33–35). Reporter mRNA is carried into the ribosome pellet through a sucrose cushion if the ribosome is puromycin-resistant and stays bound to the mRNA. As expected, the control reaction with nLuc mRNA and control recombinant Tag protein, which does not inhibit translation (9), was sensitive to puromycin as indicated by the recovery of only ∼50% of reporter mRNA in the ribosome pellet (**Figure 1C**). As previously shown (9), WT NT-hFMRP did cause puromycin-resistant mRNA•ribosome complexes (**Figure 1C**). Using published point and deletion mutants (9), we now confirm that the ncRBD is essential and sufficient to not only inhibit translation but also to form puromycin-resistant mRNA•ribosome complexes (**Figure 1C**). Noticeably, two rare FXS mutations, I304N (missense mutation in the KH2 domain) and ΔRGG+CTD (a guanine insertion within the RGG box causing a frameshift and truncated isoform), that we previously shown to have different effects on translational repression (9), have the same dichotomous effects in this assay. I304N NT-hFMRP inhibits translation and causes puromycin-resistant mRNA•ribosome complexes (**Figure 1**). Conversely, ΔRGG+CTD NT-hFRMP does not inhibit translation and causes puromycin-sensitive mRNA•ribosome complexes (**Figure 1**).

**Figure 1.**
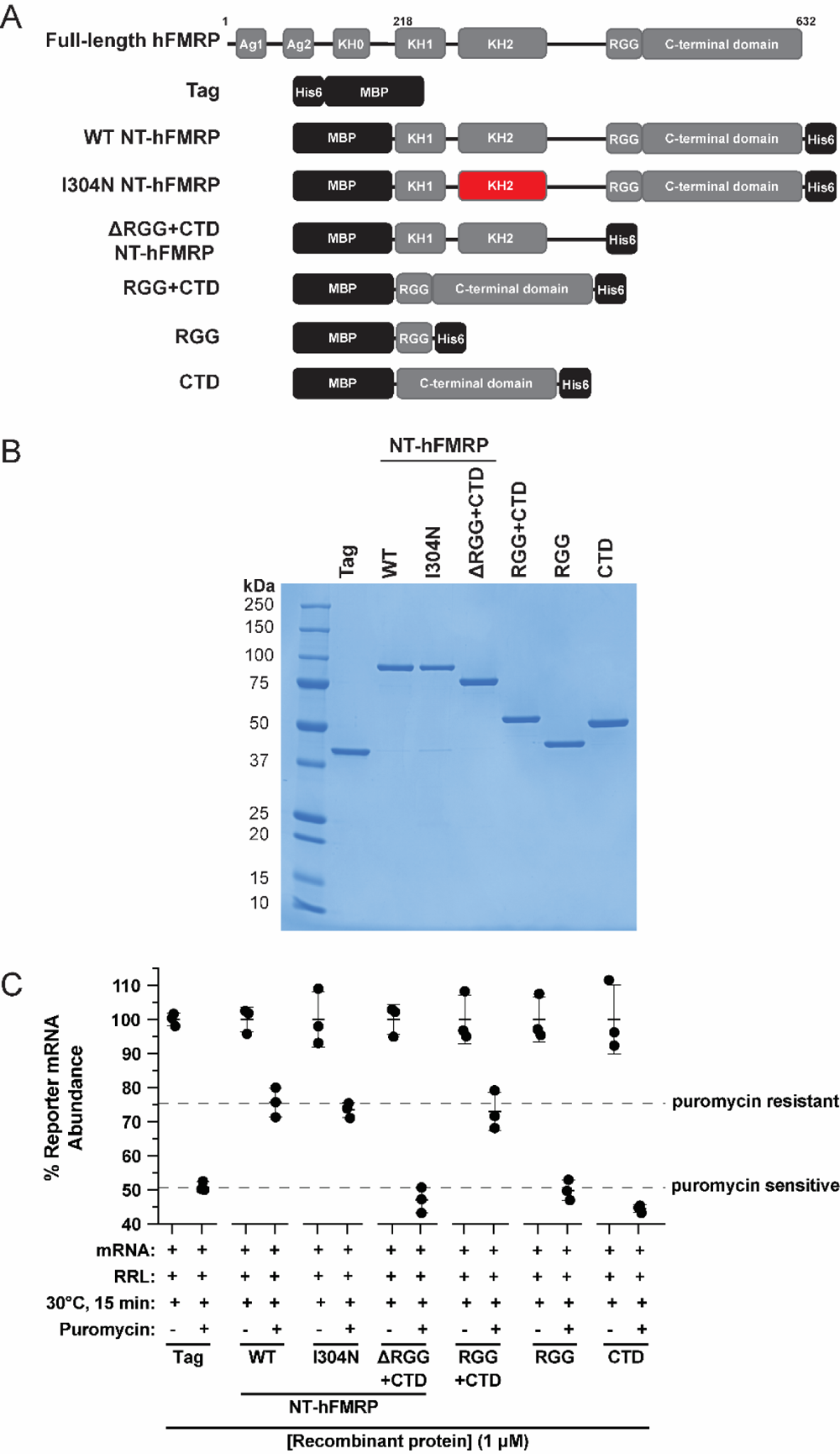
The ncRBD is essential and sufficient to cause puromycin-resistant mRNA•ribosome complexes *in vitro*. A) Schematic of full-length human FMRP and recombinant N-terminally truncated human FMRP. Mutated/truncated domains are highlighted in red. B) Coomassie stain of recombinant proteins. C) Relative quantification of nLuc reporter mRNA pelleted through a 35% (w/v) sucrose cushion after a low-speed centrifugation. nLuc•recombinant protein mRNPs were translated in RRL and treated with 0.1 mM puromycin (final) before being overlayed on the cushion and low-speed centrifugation. Final concentration of recombinant protein was 1 µM. Data are shown as mean ± SD. n=3 biological replicates. Comparisons of the puromycin treated samples were made using a two-tailed unpaired t test with Welch’s correction. Exact p-values are listed in Table S1.

### FMRP-mediated stalled ribosomes are distinct from mRNA structure-mediated stalled ribosomes

It is widely believed that stalled ribosomes often cause ribosome collisions, which in turn can stimulate NGD. One method used in the field to drive robust formation of stalled and collided ribosomes on reporter mRNAs is to delete the stop codon and insert a large GC-rich hairpin downstream (**Figure 2A**). Collided ribosomes can then be detected by treating lysates with an endonuclease (e.g., S7/micrococcal nuclease) and subsequent separation on sucrose gradients. The closely packed collided ribosomes occlude the nuclease from digesting the mRNA between ribosomes, enabling detection of stable collided ribosomes. Indeed, when compared to the control reporter mRNA, deletion of the stop codon and insertion of the hairpin in the collision reporter mRNA did result in nuclease-resistant disomes and trisomes in RRL (**Figure 2B-E and Figure S2**).

**Figure 2.**
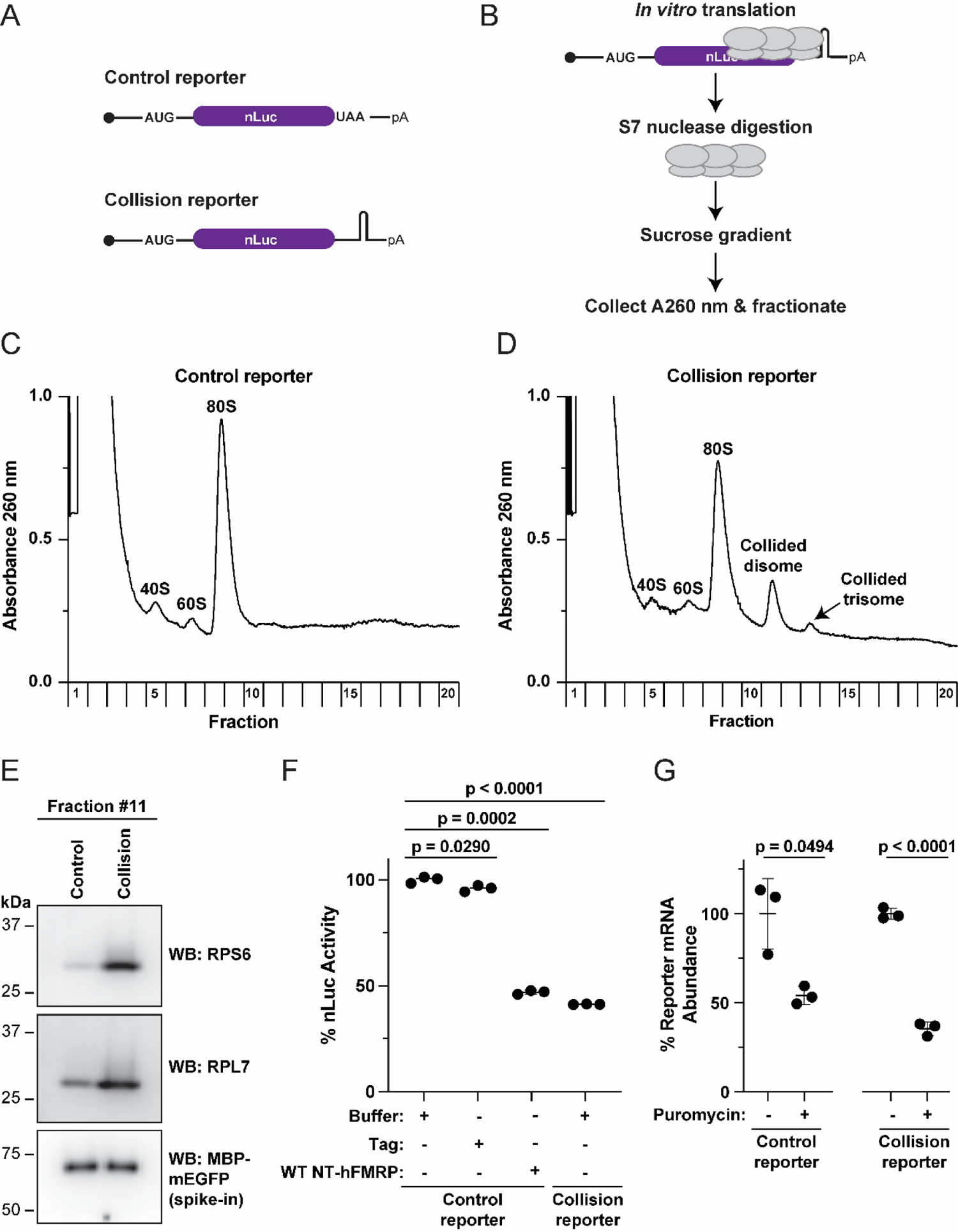
Ribosomes stalled due to mRNA structure are puromycin-sensitive. A) Schematic of control and collision reporter mRNAs. The control reporter harbors a stop codon, and the collision reporter lacks a stop codon but harbors a large GC-rich hairpin downstream of the nLuc coding sequence. B) Schematic of workflow to detect S7/micrococcal nuclease-resistant ribosome collisions on reporter mRNAs in RRL. C-D) Polysome analysis of translated control (C) and collision reporter mRNAs (D) with nuclease treatment. The collision reporter mRNA generates nuclease-resistant disomes and trisomes, with a concurrent decrease in 80S monosomes as compared to the control reporter mRNA. E) Anti-RPS6 and anti-RPL7 Western blots of fraction #11 (disome peak) which contains nuclease-resistant disomes. Recombinant MBP-mEGFP was spiked in and used as a loading control. F) *In vitro* translation of control and collision reporter mRNAs pre-incubated with protein storage buffer (Buffer) or the indicated recombinant protein (1 µM final). G) Relative quantification of nLuc control and collision reporter mRNAs pelleted through a 35% (w/v) sucrose cushion after a low-speed centrifugation. mRNAs were translated in RRL and treated with 0.1 mM puromycin (final) before being overlayed on the cushion and low-speed centrifugation. Data are shown as mean ± SD. n=3 biological replicates. Comparisons were made using a two-tailed unpaired t test with Welch’s correction.

Consistent with its inability to stimulate termination and subsequent formation of collided ribosomes, the collision reporter mRNA was translated ∼3-fold less compared to the control reporter mRNA (**Figure 2F, lanes 1 vs. 3**). A similar decrease in translation was seen with WT-hFMRP inhibition on the control reporter mRNA (**Figure 2F, lanes 2 vs. 4)**. Using the same puromycin-induced mRNA•ribosome dissociation assay as described above, we next compared ribosomes inhibited by FMRP on the control reporter mRNA to ribosomes that were stalled by the hairpin on the collision reporter mRNA. Opposite to what was observed for NT-hFMRP-mediated inhibition of the control reporter (**Figure 1C**), the mRNA•ribosome complexes formed on the collision reporter mRNA were sensitive to puromycin (**Figure 2G, right)**, which was similar to what was observed for mRNA•ribosome complexes formed on the control reporter mRNA (**Figure 2G, left)**. It should be noted that *bona fide* inhibited and stalled ribosomes via emetine treatment are hypersensitive to puromycin (36–38). This is most likely due to the ribosome being in a state that more optimally accepts puromycin as a substrate. Together, these data suggest that hFMRP-mediated stalled ribosomes are distinct from mRNA structure-mediated stalled ribosomes.

### FMRP levels regulate the stability of FMRP-targeted transcripts in cells

Despite demonstrating that recombinant NT-hFMRP inhibits translation, causes the accumulation of ribosomes on translationally repressed transcripts, and forms puromycin-resistant mRNA•ribosome complexes in RRL (9) (**Figure 1**), we were unable to detect nuclease-resistant collided ribosomes (e.g., disomes) in inhibited translation reactions using both nLuc and Firefly Luciferase (FFLuc) reporter mRNAs across multiple concentrations of FMRP **(Figure S3-S6)**. Therefore, we asked if endogenous FMRP co-sediments with nuclease-resistant collided ribosomes in Neuro2A (N2A) cells, as was shown in brain lysates (7). S7 nuclease treatment did collapse polysomes into monosomes **(Figure S7A, B and Figure S8)** and shifted signal for ribosomal proteins S6 and L7 from polysomes to monosomes on subsequent Western blots **(Figure S7C, D)**. When probing the fraction that corresponds to collided disomes (i.e., fraction #11), endogenous FMRP, RPS6, and RPL7 were significantly increased in the disome fraction in the nuclease treated samples compared to the control (non-nuclease treated) samples. This is consistent with FMRP inhibiting elongating ribosomes and causing them to collide. Importantly, this pattern was specific to ribosomal proteins and FMRP; Poly(A)-binding protein (PABPC1) was depleted from the collided disome fraction upon nuclease treatment as expected since it should not be enriched on collided ribosomes **(Figure S7E-G).**

We next asked if FMRP could stimulate NGD of its target transcripts in N2A cells. We rationalized that if FMRP does have this function, depletion of FMRP should increase steady-state levels of its NGD-regulated transcripts. Depletion of the early and essential NGD factor, Zfp598 (ZNF598 in human cells, Hel2 in yeast), should cause an increase in all *bona fide* NGD targets. Thus, the overlapping transcripts that increase would be putative FMRP-mediated NGD substrates. To test this hypothesis, we performed RNA-seq of cells after 72 hr knockdown (KD) with two independent Scramble control siRNAs, two independent siRNAs targeting *Fmr1*, or two independent siRNAs targeting *Zfp598* (**Figure 3A**). To increase the statistical power of the analysis and gain higher confidence, we performed KD in triplicate per independent siRNA and then pooled the RNA-seq data for each KD condition (i.e., Scramble KD, *Fmr1* KD, and *Zfp598* KD), for a combined n=6 per condition (**Figure 3B, C**). As expected, and in agreement with Western analysis (**Figure 3A**), each target was significantly depleted in their respective KD samples (**Figure 3B, C**, blue data points). Using a two-fold RNA fold change cut off (adjusted p<0.05), we identified 132 and 55 transcripts that increased with *Fmr1* and *Zfp598* depletion, respectively (**Figure 3B, C**, Table S2, Table S3). Of these transcripts, 16 increased at least 2-fold in both KD conditions compared to the Scramble controls (**Figure 3B, C** red data points and **Figure 3D**), representing putative FMRP-mediated NGD substrates. Upon mining published data sets, we found that 14 of the 16 putative FMRP-mediated NGD substrates have been identified in a mouse brain FMRP crosslinking immunoprecipitation (CLIP)-seq data set (**Figure 3E** and Table S4) (39), suggesting that the observed effects are direct rather than secondary or downstream.

**Figure 3.**
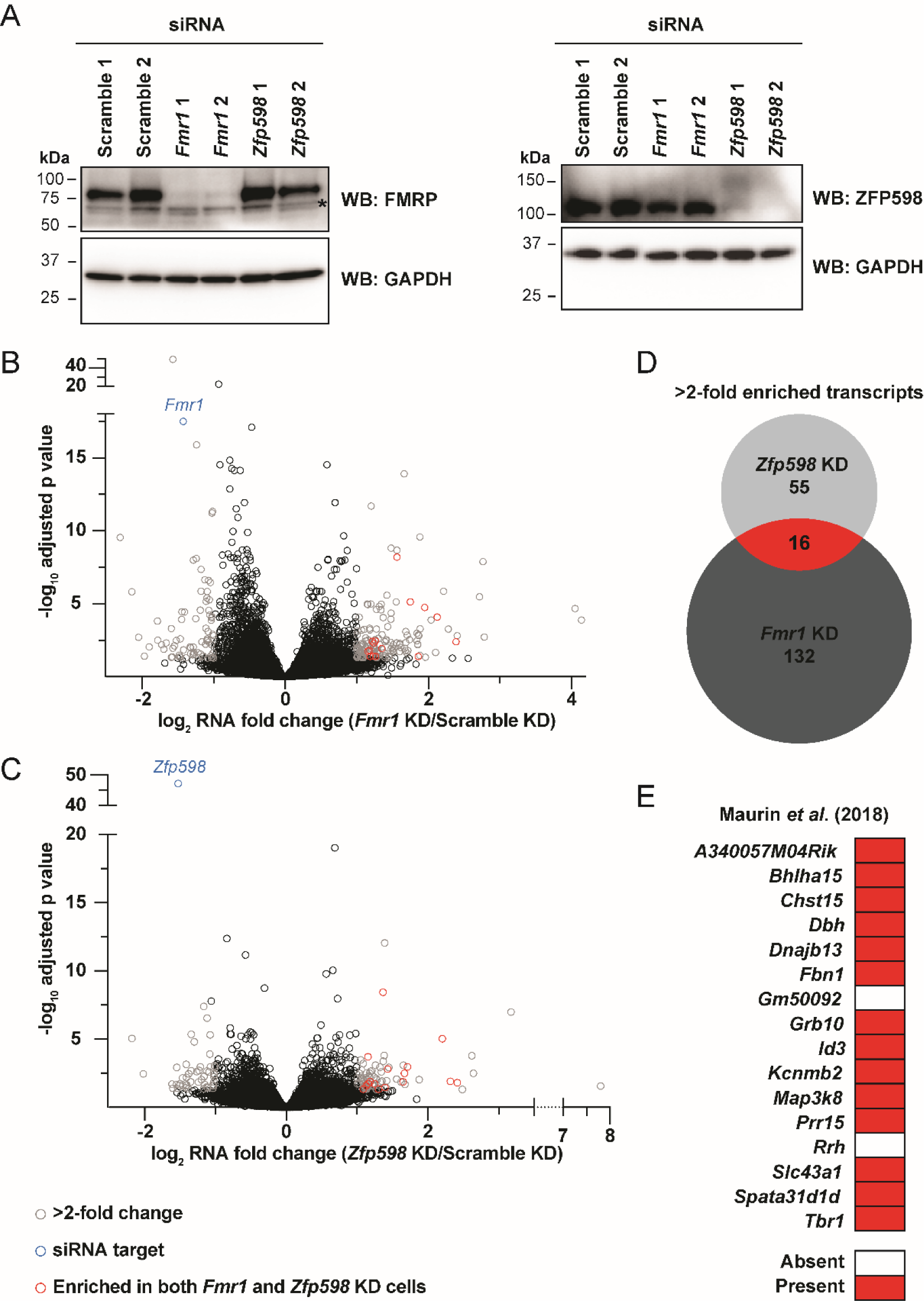
RNA-seq reveals 16 transcripts as putative FMRP-mediated NGD targets in mouse N2A cells. A) Anti-FMRP and anti-ZFP98 Western blots of N2A cell lysates treated with the indicated siRNAs. GAPDH was used as a loading control. *Denotes non-specific immunoreactivity. B-C) Volcano plots of RNA-seq data comparing Scramble control to *Fmr1* KD (B) or *Znf598* KD (C). Two independent siRNAs were used in triplicate per target for n=6 per target. Transcripts enriched >2-fold (adjusted p<0.05) in both *Fmr1* and *Zfp598* KD conditions are highlighted in red. D) Venn diagram of >2-fold enriched transcripts in *Fmr1* and *Zfp598* KD conditions. 16 transcripts are enriched in both KD conditions tested (red). E) 14 of the16 identified transcripts (red in D) are found in mouse brain FMRP CLIP-seq in Maurin *et al*. (39).

To validate these results, we pursued measuring the change in mRNA half-life (t_1/2_) in N2A cells after *Fmr1* or *Zfp598* depletion using Roadblock-qPCR, which takes advantage of 4-thiouridine (4SU) incorporation into mRNA and the ability to form 4SU-*N*-ethylmaleimide (NEM) adducts that block reverse transcriptase (40). Using the same approach as above, two independent siRNAs were used per KD target in triplicate, for a combined n=6, per condition per timepoint over an 8 hr time course. 72 hrs after KD, N2A cells were treated with 4SU and total RNA was harvest at the indicated timepoints. In agreement with Western analysis and RNA-seq (**Figure 4A-C**), at t=0 hr, *Fmr1* and *Zfp598* mRNAs were significantly depleted in their respective KD samples (**Figure 4A**, B). Validating the Roadblock-qPCR approach in these cells, we did observe the robust stability of *Actb* mRNA (**Figure 4C**) and instability of *Sesn2* mRNA (**Figure 4D**, black data points and curve) over 8 hrs with the Scramble negative control siRNAs. Importantly, the short t_1/2_ of *Sesn2* mRNA was not sensitive to *Fmr1* KD nor *Zfp598* KD when comparing the t_1/2_ 95% CI (**Figure 4D, E**). Importantly, we did observe a ∼1.7-fold increase that was statistically significant in the t_1/2_ of the FMRP-targeted *Id3* mRNA with both *Fmr1* and *Zfp598* KD. However, *Id3* mRNA stability was not fully rescued, suggesting it is naturally unstable and/or under regulation of other mRNA decay pathways. Similar increased stability was observed for *Dbh*, *Map3k8*, and *Tbr1* mRNAs (Figure S9), all of which were identified in our RNA-seq analysis as putative FMRP-mediated NGD substrates and were identified in a mouse brain FMRP CLIP-seq dataset (39). In total, these data provide evidence that FMRP can stall ribosomes to stimulate NGD on targeted transcripts.

**Figure 4.**
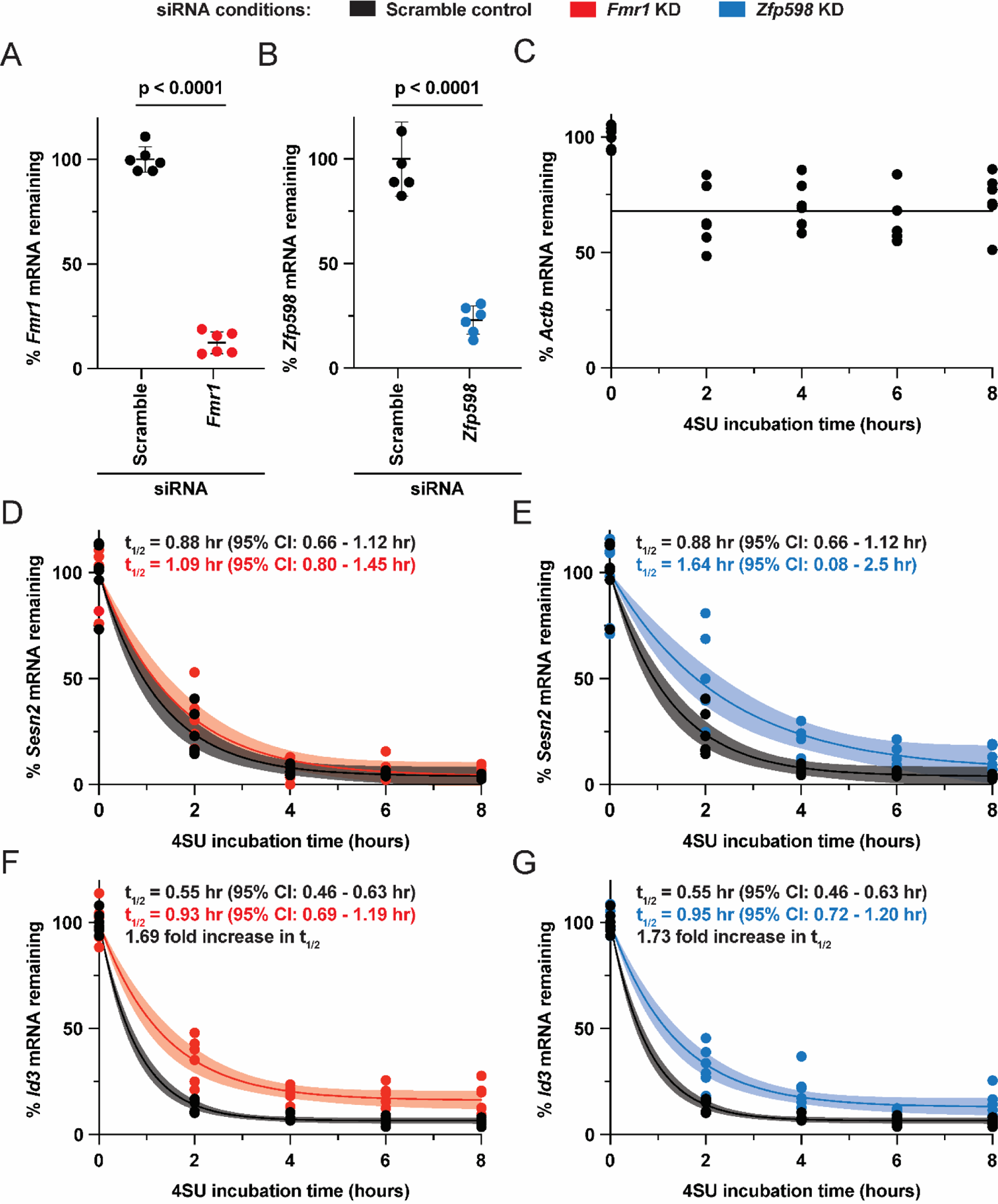
Depletion of Fmr1 and the key NGD factor Zfp598 equally stabilize *Id3* mRNA in N2A cells. A, B) RT-qPCR was used to confirm depletion of *Fmr1* (A) and *Zfp598* (B) mRNA 72 hrs after KD at t=0 hr 4SU incorporation. Data are shown as the mean ± SD. n=6 biological replicates. Comparisons were made using a two-tailed unpaired t test with Welch’s correction. C-G) Roadblock-qPCR was used to measure mRNA half-lives (t_1/2_) of *Actb* mRNA (Scramble negative control) (C), as well as *Sesn2* mRNA (D-E) and *Id3* mRNA (F-G). The Scramble negative control is in black (and is the same in D & E, as well as in F & G), *Fmr1* KD is in red, and *Zfp598* KD is in blue. n=6 biological replicates. mRNA t_1/2_ was determined by one phase decay trend lines calculated by nonlinear regression. The 95% confidence interval range is reported and is shown as a watermark.

## DISCUSSION

Data presented here add to the existing evidence that stalled ribosomes can have different properties depending on the effector. For example, emetine and cycloheximide bind to the E site of the 40S and 60S subunits of the 80S ribosome, respectively. Both inhibit and stall elongating ribosomes (41, 42) yet they have opposite effects on puromycylation (36–38, 42), where emetine enhances and cycloheximide inhibits puromycylation (36–38, 42). It is thought that emetine traps ribosomes in a state that more optimally accepts puromycin as a substrate (36–38). Here we show that ribosomes inhibited by FMRP and ribosomes inhibited by mRNA structure are distinct as only the former are puromycin-resistant.

High doses of translation elongation inhibitors stall most ribosomes preventing collisions, whereas intermediate doses cause a select few to act as roadblocks for trailing ribosomes (26, 43–45), ultimately causing ribosome collisions and NGD. Analogous to intermediate doses, in mining published datasets we found that the 16 putative FMRP-mediated NGD substrates represent low-density FMRP targets (39). High-density FMRP targets from the same and other FMRP CLIP-seq datasets (7, 39, 46–52), many of which are expressed in N2A cells, were not identified in our experiments as being regulated by NGD, suggesting that stimulating NGD is a minor function of FMRP. It is also possible that a high-density of bound FMRP inhibits most ribosomes and aids in preventing ribosome collisions. Furthermore, some data suggest that in neurons, most mRNAs are translated by monosomes (53), where ribosome collisions cannot be formed. Concordantly, neurons may have optimized their translation status to avoid causing widespread FMRP-mediated ribosome collisions and prevent mRNA instability in order to enable rapid regulated local translation in response to stimulation (54, 55).

The 16 putative FMRP-mediated NGD substrates encode several neuronal proteins **(Table S4)**, many of which could contribute to documented FXS phenotypes (56), including those involved in neurotransmission (*Dbh*), cell differentiation regulation in the hippocampus (*Id3*), neuronal excitability (*Kcnmb2*), as well as neuronal migration and axonal projection (*Tbr1*). Additionally, *Prr15* is known to be involved in glioblastoma, and *Rrh* is a retinal-associated gene. Notably, the Darnell group identified *Id3* as an FMRP target over two decades ago (57). ID3 is expressed in the hippocampus (58), an affected brain area in FXS (59), and binds to and inhibits the basic helix-loop-helix transcription factors, consequently inhibiting cell differentiation. Our data suggest that FMRP normally controls *Id3* mRNA stability. Thus, when FMRP is lost in FXS, the increased *Id3* mRNA levels could cause a concatenate increase in ID3 protein in the proliferative region of the hippocampus, resulting in aberrant inhibition of granule cell and dentate precursor cell differentiation (58).

## EXPERIMENTAL PROCEDURES

Full experimental procedures can be found in the Supporting Information. Exact p-values for Figure 1 can be found in Table S1. RNA-seq data analysis can be found in Table S2 and Table S3. Reported gene functions for the hits in Figure 4 can be found in Table S4. Silencer Select siRNAs (Thermo) used in this study can be found in Table S5. Oligonucleotides used in this study can be found in Table S6.

## DATA AVAILABILITY

All data is within the article or in the Supporting Information except for the raw RNA-seq data which has been deposited to the NCBI GEO under ascension number GSE254586.

## ACKNOWLEDGMENTS

We thank members of the Kearse lab, Karin Musier-Forsyth, and Kurt Fredrick for input and thoughtful discussions.

## AUTHOR CONTRIBUTIONS

M. R. S. and M. G. K. conceptualization; M.R.S, B.P., W.T., and M.G.K investigation; M.R.S. and M.G.K. writing–original draft; M.R.S, B.P., W.T., and M.G.K writing–review & editing; W.T. and M. G. K. supervision.

## FUNDING AND ADDITIONAL INFORMATION

M. R. S. was supported by The Ohio State University Fellowship and The Ohio State University Center for RNA Biology Graduate Fellowship. B.P. was supported by The Ohio State University Center for RNA Biology Graduate Fellowship and The Ohio State University Presidential Fellowship. This work was supported by NIH grant R35GM142580 to W. T. and NIH grant R35GM146924 to M.G. K. The Ohio State University Comprehensive Cancer Center Genomics Shared Resource (OSUCCC GSR) is supported by NIH grant P30CA016058. The content is solely the responsibility of the authors and does not necessarily represent the official views of the National Institutes of Health.

## SUPPORTING INFORMATION

### SUPPLEMENTAL EXPERIMENTAL PROCEDURES

#### Plasmids

We have previously described the design of a G-quadruplex lacking nLuc coding sequence, that was based off of pNL1.1 (Promega), harboring the human β-globin 5ʹ UTR that was synthesized by Integrated DNA Technologies and cloned into pcDNA3.1(+) (9). To generate a reporter that robustly stimulates ribosome collisions, the stop codon of nLuc was removed and a large GC-rich hairpin (HP) was inserted downstream using the Q5 Site-Directed Mutagenesis Kit (NEB # E0552S). pCRII/FFLuc, which contains the FFLuc coding sequence from pGL4.13 (Promega) downstream from the T7 RNA polymerase promoter, was previously described (Kearse *et al*, 2016).

An *E. coli* optimized coding sequence for human FMRP (isoform 1) was designed and synthesized by Genscript, and then subcloned into pET His6 MBP TEV LIC cloning vector (1M), which was a gift from Scott Gradia (Addgene plasmid # 29656), through ligation-independent cloning (LIC) using Novagen’s LIC-qualified T4 DNA polymerase (Sigma # 70099-M) as described by Q3 Macrolab (http://qb3.berkeley.edu/macrolab/). The His6-tag was deleted from the N-terminus and inserted at the C-terminus. The NT-hFMRP sequence included a P451S mutation to prevent ribosome stalling at a poly-proline stretch and formation of truncated recombinant protein, as previously described (32). Point mutations and deletions were achieved using the Q5 Site-Directed Mutagenesis Kit. To be as consistent as possible across the previous literature, we refer to the RGG box motif as to the minimal region identified by Darnell and colleagues that bound Sc1 RNA with highest affinity (57).

All plasmids were propagated in TOP10 *E. coli* (Thermo Fisher # C404006), purified using the PureYield Plasmid Miniprep or Midiprep Systems (Promega # A1222 and A2495), and validated by Sanger sequencing at The Ohio State University Comprehensive Cancer Center Genomics Shared Resource (OSUCCC GSR). Sequences of the reporters and recombinant proteins are provided in the Supporting Information.

#### Reporter mRNA *in vitro* transcription

All nLuc plasmids were linearized with XbaI and purified using a Zymo DNA Clean & Concentrator 25 Kit (Zymo Research # D4065). pCRII/FFLuc was linearized with HindIII using a Zymo DNA Clean & Concentrator 25 Kit. DNA was transcribed into mRNA which was co-transcriptionally capped with the Anti-Reverse Cap Analog (ARCA) 3′-O-Me-m7G(5′)ppp(5′)G (NEB # S1411) using the HiScribe T7 High Yield RNA Synthesis Kit (NEB # E2040). Our standard 10 µL reactions used 0.5 µg of linear plasmid template and an 8:1 ARCA:GTP ratio. Reactions were incubated at 30°C for 2 hrs, then incubated with 20 U of DNaseI (NEB # M0303S) at 37°C for 15 min, and then purified using a Zymo RNA Clean & Concentrator 25 Kit (Zymo Research # R1018). Reporter mRNA was eluted in 75 µL of RNase-free water, aliquoted in single use volumes, and stored at -80°C. Reporter mRNA integrity was confirmed by denaturing formaldehyde agarose gel electrophoresis and ethidium bromide visualization. We routinely found the 30°C incubation resulted in less observable truncated products than incubation at 37°C and did not significantly affect yield for our purposes. mRNA translation in nuclease-treated rabbit reticulocyte lysate (RRL) is poly(A)-independent (Soto Rifo *et al*., 2007); thus, we omitted polyadenylating reporter mRNAs in this study.

#### Recombinant protein expression and purification

All recombinant proteins were expressed in Rosetta 2(DE3) *E. coli* (Sigma # 71397-4) using MagicMedia *E. coli* Expression Medium (Thermo Fisher # K6803) supplemented with 50 µg/mL kanamycin and 35 µg/mL chloramphenicol for auto-induction. A 5 mL starter culture in LB media supplemented with 50 µg/mL kanamycin, 35 µg/mL chloramphenicol, and 1% glucose (w/v) was inoculated with a single colony and grown overnight at 37°C, 250 rpm. 1 mL of a warm and fresh overnight starter culture was then used to inoculate 50 mL of room temperature MagicMedia and incubated for 48-72 hrs at 18°C, 160 rpm in a 250 mL baffled flask. After auto-induction, cultures were pelleted and stored at -20°C for purification later. Recombinant proteins were purified using a dual affinity approach, first using the C-terminal His6-tag, then the N-terminal MBP-tag. Cell pellets were resuspended and lysed with BugBuster Master Mix (Sigma # 71456) using the recommended 5 mL per 1 g wet cell pellet ratio for 10 min at room temperature with gentle end-over-end rotation (10-15 rpm). Lysates were placed on ice and kept cold moving forward. Lysates were cleared by centrifugation for 20 min at 18,000 rcf in a chilled centrifuge (4°C). Lysates were then incubated with HisPur Cobalt Resin (Thermo Fisher # 89965) in a Peirce centrifugation column (Thermo # 89897) for 30 min at 4°C with gentle end-over-end rotation. Columns were centrifuged in a pre-chilled (4°C) Eppendorf 5810R for 2 min at 700 rcf to eliminate the flow through and then were washed 5X with two resin-bed volumes of ice-cold Cobalt IMAC Wash Buffer (50 mM Na3PO4, 300 mM NaCl, 10 mM imidazole; pH 7.4) in a pre-chilled (4°C) Eppendorf 5810R for 2 min at 700 rcf. His-tagged proteins were then eluted in a single elution step with two resin-bed volumes of ice-cold Cobalt IMAC Elution Buffer (50 mM Na3PO4, 300 mM NaCl, 150 mM imidazole; pH 7.4) by gravity flow. Eluates were then incubated with Amylose resin (NEB # E8021) in a centrifugation column for 2 hrs at 4°C with gentle end-over-end rotation (10-15 rpm). Columns were washed 5X with at least two bed-volumes of ice-cold MBP Wash Buffer (20 mM Tris-HCl, 200 mM NaCl, 1 mM EDTA; pH 7.4) by gravity flow. MBP-tagged proteins were then eluted by a single elution step with two resin-bed volumes of ice-cold MBP Elution Buffer (20 mM Tris-HCl, 200 mM NaCl, 1 mM EDTA, 10 mM maltose; pH 7.4) by gravity flow. Recombinant proteins were then desalted and buffer exchanged into Protein Storage Buffer (25 mM Tris-HCl, 125 mM KCl, 10% glycerol; pH 7.4) using a 7K MWCO Zeba Spin Desalting Column (Thermo Fisher # 89892) and, if needed, concentrated using 10K MWCO Amicon Ultra-4 (EMD Millipore # UFC803024). Recombinant protein concentration was determined by Pierce Detergent Compatible Bradford Assay Kit (Thermo Fisher # 23246) with BSA standards diluted in Protein Storage Buffer as well as SDS-PAGE and Coomassie staining before aliquoting in single use volumes, snap freezing in liquid nitrogen, and storage at -80°C.

#### mRNP formation and *in vitro* translation

*In vitro* translation was performed in the dynamic linear range as previously described but adapted to translate mRNPs (9, 23, and Kearse *et al*., 2016). 30 nM *in vitro* transcribed nLuc reporter mRNA was diluted in RNA Dilution Buffer (10 mM Tris-HCl, 5 mM Mg(OAc)_2_, 100 mM KCl; pH 7.4). In a total of 4 µL, 30 fmol of nLuc reporter mRNA was mixed with 0-10 picomol of recombinant protein and 100 picomol of UltraPure BSA (Thermo Fisher # AM2618) on ice for 1 hr. UltraPure BSA stock was diluted in protein storage buffer and its addition was necessary to prevent non-specific binding of the reporter mRNA to the tube. For *in vitro* translation reactions, 6 µL of a Rabbit Reticulocyte Lysate (RRL) master mix was added to each 4µL mRNP complex. 10 μL *in vitro* translation reactions were performed in the linear range using 3 nM mRNA in the Flexi RRL System (Promega # L4540) with final concentrations of reagents at 30% RRL, 10 μM amino acid mix minus leucine, 10 μM amino acid mix minus Methionine, 0.5 mM Mg(OAc)_2_, 100 mM KCl, 8 U murine RNase inhibitor (NEB # M0314), 0-1 µM recombinant protein, and 10 µM UltraPure BSA. Reactions were incubated for 30 min at 30°C, terminated by incubation on ice and diluted 1:5 in Glo Lysis Buffer (Promega # E2661). 25 μL of prepared Nano-Glo reagent (Promega # N1120) was mixed with 25 μL of diluted reaction and incubated at room temperature for 5 min in the dark (with gentle shaking during the first minute), and then read on a Promega GloMax Discover Multimode Microplate Reader. FFLuc mRNA was treated and translated exactly the same; FFLuc luminescence was measured exactly the same but used ONE-Glo (Promega # E6110) instead of Nano-Glo.

#### mRNA•ribosome dissociation assays with puromycin

We have previously described in detail and in a complete methods manuscript the use of the ability of puromycin to dissociate ribosomes from reporter mRNA with a low-speed sucrose cushion (9, 23). nLuc reporter mRNA translation was performed as described above except that translation was limited to 15 min at 30°C. Samples were then placed on ice for 3 min before the addition of 0.1 mM puromycin (final) and further incubation at 30°C for 30 min. Control samples lacking puromycin (water added instead) were kept on ice. Cycloheximide (1.43 mg/mL final) was then added to all samples to preserve ribosome complexes on mRNAs and halt puromycin incorporation. In a separate tube, FFLuc reporter mRNA was translated as described above (3 nM mRNA conditions) for 15 min at 30°C and was terminated by the addition of 1.43 mg/mL cycloheximide (final) and incubation on ice.

The treated nLuc and FFLuc translation reactions (from above) were combined on ice and then mixed with an equal volume (28 µL) of ice-cold 2X Ribosome Dilution Buffer (40 mM Tris-HCl, 280 mM KCl, 20 mM MgCl_2_, 200 μg/ml cycloheximide, 2 mM DTT; pH 7.4). The entire 56 μl volume was then layered on top of 130 μl of ice-cold 35% (w/v) buffered sucrose (20 mM Tris-HCl, 140 mM KCl, 10 mM MgCl_2_, 100 μg/mL cycloheximide, 1 mM DTT; pH 7.4) in a pre-chilled 7 mm x 20 mm thick-walled polycarbonate ultracentrifuge tubes (Thermo Scientific # 45233) and centrifuged in a S100AT3 rotor at 4°C for 60 min at 50,000 x g (43,000 rpm) in a Sorvall Discovery M120 SE Micro-Ultracentrifuge. The supernatant was then discarded and each pellet was resuspended in 0.5 mL of TRIzol (Thermo Fisher # 15596018). Total RNA was extracted from each pellet following the manufacturer’s protocol with glycogen (Thermo Fisher # R0561) added at the isopropanol precipitation step. The resulting RNA pellet was resuspended in 30 μL nuclease-free water. 16 μL of extracted RNA was converted to cDNA using iScript Reverse Transcription Supermix for RT-qPCR (Bio-Rad # 1708841). cDNA reactions were then diluted 10-fold with nuclease-free water and stored at -20°C or used immediately. RT-qPCR was performed in 15 μL reactions using iTaq Universal SYBR Green Supermix (Bio-Rad # 1725124) in a Bio-Rad CFX Connect Real-Time PCR Detection System with 1.5 μL diluted cDNA and 250 nM (final concentration) primers. nLuc reporter mRNA abundance was normalized to the spiked-in control FFLuc mRNA using the Bio-Rad CFX Maestro software (ΔΔCt method). Abundance of total signal was calculated using *Q_n_ = 2^ΔΔCt^* and *P = 100 × Q_n_/Q_total_* as previously described (Pringle *et al*., 2019). Primers for RT-qPCR can be found in **Table S6**.

#### Cell culture and siRNA knockdowns

N2A cells were obtained from ATCC (# CCL-131) and maintained in high glucose DMEM (Thermo # 11995065) supplemented with 10% heat-inactivated FBS and 1% penicillin-streptomycin in standard tissue culture-treated plastics at 37°C with 5% CO_2_. N2A cells were seeded in 12-well plates 24 hrs prior to knockdown, and then transfected with Silencer Select siRNAs (Thermo) using Lipofectamine RNAiMAX (Thermo # 13778150) following the manufacturer’s recommendation. At the time of transfection, N2A cells were at ∼20% confluency. 24 hrs post siRNA transfection, the media was changed. After 72 hr transfection, total RNA or protein was harvested by TRIzol or RIPA buffer, respectively. Silencer Select siRNAs used in this study are listed in **Table S5**.

#### Sucrose gradient ultracentrifugation

*In vitro* translation reactions were scaled up 10-fold to 100 µL. After 30 min at 30°C, reactions were transferred to ice, 1 µL of 100 mg/mL cycloheximide was added (∼1 mg/mL final), and samples were snap frozen in liquid nitrogen and stored at -80°C. When ready to perform sucrose gradients, samples were then thawed on ice, 5 µL of 10 mM CaCl_2_ (∼0.5 mM final) and 5.3 µL of 3,000 U/mL S7 micrococcal nuclease (Thermo # EN0181; stock at 30,000 U/mL in PBS; ∼150 U/mL final) was added. After nuclease digestion at 25°C for 10 min, samples were quenched by adding 2.2 µL of 50 mM EGTA (∼1 mM final). Samples were then diluted with an equal volume of ice-cold 2X Polysome Dilution Buffer (40 mM Tris-HCl, 280 mM KCl, 20 mM MgCl_2_, 2 mM DTT, 200 µg/mL cycloheximide; pH 7.4), gently mixed, and layered on top of a linear 10-50% (w/v) buffered sucrose gradient (20 mM Tris-HCl, 140 mM KCl, 10 mM MgCl_2_, 1 mM DTT, 100 μg/mL cycloheximide; pH 7.4) in a 14 mm × 89 mm thin-wall Ultra-Clear tube (Beckman # 344059) that was formed using a Biocomp Gradient Master. Gradients were centrifuged at 35K rpm for 120 min at 4°C in a SW-41Ti rotor (Beckman) with maximum acceleration and no brake using a Beckman Optima L-90 Ultracentrifuge. Gradients were subsequently fractionated into 0.5 mL volumes using a Biocomp piston fractionator with a TRIAX flow cell (Biocomp) recording a continuous A_260 nm_ trace.

Per replicate, N2A cells were seeded in four 15 cm tissue culture treated plates and allowed to grow for 2-3 days until ∼70% confluent. Cells were briefly treated with 100 µL/mL cycloheximide for 2 min, then rinsed with ice-cold PBS with 100 µL/mL cycloheximide, harvested using a cell lifter, pooled, pelleted at 500 rcf for 5 min at 4°C in a pre-chilled centrifuge, snap froze in liquid nitrogen, and stored at -80°C. Cell pellets were thawed on ice and then lysed in 950 μL ice-cold Polysome Lysis Buffer (20 mM Tris-HCl, 140 mM KCl, 10 mM MgCl_2_, 1 mM DTT, 100 μg/mL cycloheximide, 1% (v/v) Triton X-100; pH 7.4) for 10 min on ice. Cell debris was then cleared at 13,000 rcf at 4°C in a pre-chilled microcentrifuge. The supernatant was taken and 450 µL was aliquoted into two new pre-chilled microcentrifuge tubes. 22.5 μL of 10 mM CaCl_2_ (∼0.5 mM final) and 2.4 µL of either 30,000 U/mL S7 micrococcal nuclease (∼150 U/mL final) or PBS was added. After nuclease digestion at 25°C for 10 min, samples were quenched by adding 9.5 µL of 50 mM EGTA (∼1 mM final). 400 µL was then layered on top of a linear 10-50% (w/v) buffered sucrose gradient (20 mM Tris-HCl, 140 mM KCl, 10 mM MgCl_2_, 1 mM DTT, 100 μg/mL cycloheximide; pH 7.4), centrifuged, and fractionated as described above.

#### Western blotting

When assaying sucrose gradient fractions, 100 μL of the 500 μL collected fractions was added and mixed with 2 μL of 1.4 μΜ MBP-mEGFP, which served as a spike-in loading control. 34 μL of 4X reducing SDS sample buffer (Bio-Rad # 1610737) was then added, mixed, and heated at 70°C for 15 min. 30 µL was then separated by Tris-Glycine SDS-PAGE (Thermo # XP04200BOX) and transferred on to 0.2 µm PVDF membrane (Thermo # 88520). Membranes were then blocked with 5% (w/v) non-fat dry milk in TBST (1X Tris-buffered saline with 0.1% (v/v) Tween 20) for 30 min at room temperature before overnight incubation with primary antibodies in TBST at 4°C with gentle rotation/rocking. After three 10 min washes with TBST, membranes were incubated with HRP-conjugated secondary antibody in TBST for 1 hr at room temperature and then washed again with three 10 min washes with TBST. Chemiluminescence was performed with SuperSignal West Pico PLUS (Thermo # 34577) for GFP, RPS6, RPL7, and PABP and with SuperSignal West Atto Maximum Sensitivity Substrate (Thermo # A38555) for FMRP, and imaged using an Azure Sapphire Biomolecular Imager. Rabbit anti-GFP (Cell Signaling # 2956S) was used at 1:1,000. Rabbit anti-RPS6 (Cell Signaling # 2217) was used at 1:1,000. Rabbit anti-RPL7 (abcam # ab72550) was used at 1:1,000. Rabbit anti-PABP (abcam # ab21060) was used at 1:1,000. HRP-conjugated goat anti-rabbit IgG (H+L) (Thermo # 31460) was used at 1:10,000 for GFP and PABP, 1:100,000 for FMRP, and 1:30,000 for RPS6 and RPL7.

When assaying N2A cell lysates after 72 hr knockdown, whole cell lysates were prepared using RIPA buffer. Briefly, cells were placed on a bed of ice and media was aspirated before a gentle 1 mL ice-cold PBS rinse. Cells were then lysed in 300 μL of ice-cold RIPA buffer for 10 min at 4°C with gentle rocking. The entire lysate (including any apparent cell debris) was mixed with 100 μL of 4X reducing SDS sample buffer and heated to 70°C for 15 min. Samples were then homogenized by syringing 6X through a 28G needle. 30 µL was then separated by Tris-Glycine SDS-PAGE, transferred on 0.2 µm PVDF, and probed as described above. Rabbit anti-ZNF598 (Thermo # PA5-59777) was used at 1:1,000. Rabbit anti-FMRP (Abcam # ab17722) was used at 1:1,000. Rabbit anti-GAPDH (Cell Signaling # 5174) was used at 1:1,000. HRP-conjugated goat anti-rabbit IgG (H+L) (Thermo # 31460) was used at 1:10,000 for ZNF598 and FMRP, and at 1:30,000 for GAPDH. Chemiluminescence was performed with SuperSignal West Pico PLUS and imaged using an Azure Sapphire Biomolecular Imager.

#### RNA-seq and mRNA decay analyses via Roadblock-qPCR

After 72 hr knockdown, total RNA was extracted with TRIzol reagent following the manufacturer’s protocol. RNA concentration and purity was determined by UV spectroscopy (i.e., Nanodrop). RNA library preparation (with RNA quality validation and rRNA depletion included) and sequencing was conducted by Novogene. Adapters and poorly sequenced reads were removed from raw sequencing data using TrimGalore (Kechin *et al*., 2017). The quality of sequencing reads following trimming and filtration was assessed using FastQC (Andrews, 2015). Processed sequencing reads were aligned to the mouse reference genome (GRCm39) using STAR (Dobin *et al*., 2013). Sequence coverage BigWig files were generated using STAR alignment BAM output using the deepTools tool bamCoverage, using counts per million (CPM) normalization (Ramirez *et al*., 2014). Genomic features were quantified by counting the number of reads aligning to exons of genes using FeatureCounts (Liao *et al*., 2013). Normalization and differential gene expression analysis was then preformed using the R package DESeq2 (Love *et al*., 2014). The pipeline used for the pre-processing, alignment, and post-alignment analysis of RNA-seq data can be found at https://doi.org/10.5281/zenodo.8302724. Any other custom scripts used in this analysis are available upon request. All raw RNA-seq data has been deposited in the NCBI Gene Expression Omnibus (GEO).

Roadblock-qPCR was used to determine mRNA half-lives and was performed as described by the Thoreen lab (40). 72 hr post knockdown, media was replaced with pre-warmed completed supplemented with 400 μΜ 4SU (stock at 80 mM in DMSO; Sigma #T4509-100MG). Timepoints were taken immediately (0 hr), 2, 4, 6, and 8 hrs later by aspirating media and adding 1 mL of TRIzol. After extracting total RNA by following the manufacture’s recommendations, 3 μg of RNA was modified in NEM Reaction Buffer (50 mM Tris-HCl, 1 mM EDTA, and 50 mM NEM; pH8) in a 50 μL reaction at 42°C for 90 min. 1 M NEM (Sigma # 3876) was made with 100% EtOH, aliquoted, and stored at -20°C. The reaction was quenched by addition of 20 mM DTT (final) and RNA was purified using an RNA Clean & Concentrator-5 Kit (Zymo # R1013) with a final elution volume of 15 μL. 1 μg of recovered RNA was used to generate cDNA for RT-qPCR using the iScript Reverse Transcription Supermix for RT-qPCR (Bio-Rad # 1708841). cDNA reactions were then diluted 10-fold with nuclease-free water and stored at −20°C or used immediately. RT-qPCR was performed in 15 μl reactions using iTaq Universal SYBR Green Supermix (Bio-Rad# 1725124) in a Bio-Rad CFX Connect Real-Time PCR Detection System with 1.5 μl diluted cDNA and 250 nM (final concentration) primers. Target mRNA levels were normalized to 18S rRNA, and half-lives were calculated using one phase decay trend lines calculated by nonlinear regression in GraphPad Prism 10.0.3. All data are reported and the 95% confidence interval was included as a watermark when appropriate. All RT-qPCR primer sequences are available in **Table S6**.

## SUPPLEMENTAL FIGURES

**Figure S1.**
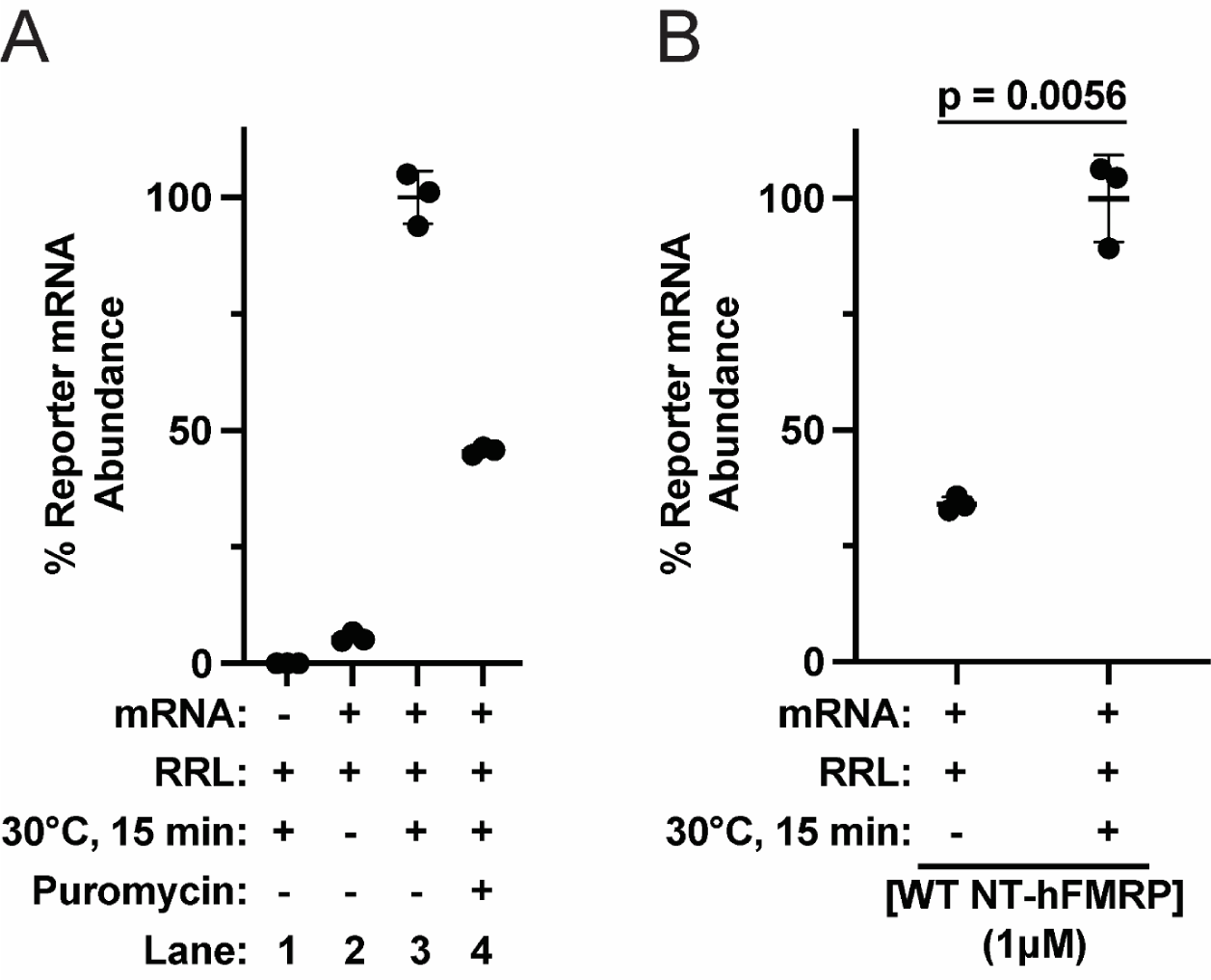
nLuc mRNA pelleting through the sucrose cushion is translation dependent. A) Relative quantification of nLuc reporter mRNA pelleted through a 35% (w/v) sucrose cushion after a low-speed centrifugation. Lane 1 is a negative control lacking nLuc mRNA. Lane 2 is a negative control containing mRNA in RRL but not incubated at 30°C to start translation. Lane 3 is nLuc mRNA in RRL translated for 15 min at 30°C. Lane 4 is nLuc mRNA in RRL translated for 15 min at 30°C and then incubated with 0.1 mM puromycin (final) for 30 min at 30°C. Data are shown as mean ± SD. n=3 biological replicates. B) Relative quantification of nLuc reporter mRNA pelleted through a 35% (w/v) sucrose cushion after a low-speed centrifugation with WT NT-hFMRP (1 µM final) with and without translation (15 min at 30°C). Data are shown as mean ± SD. n=3 biological replicates. Comparisons were made using a two-tailed unpaired t test with Welch’s correction.

**Figure S2.**
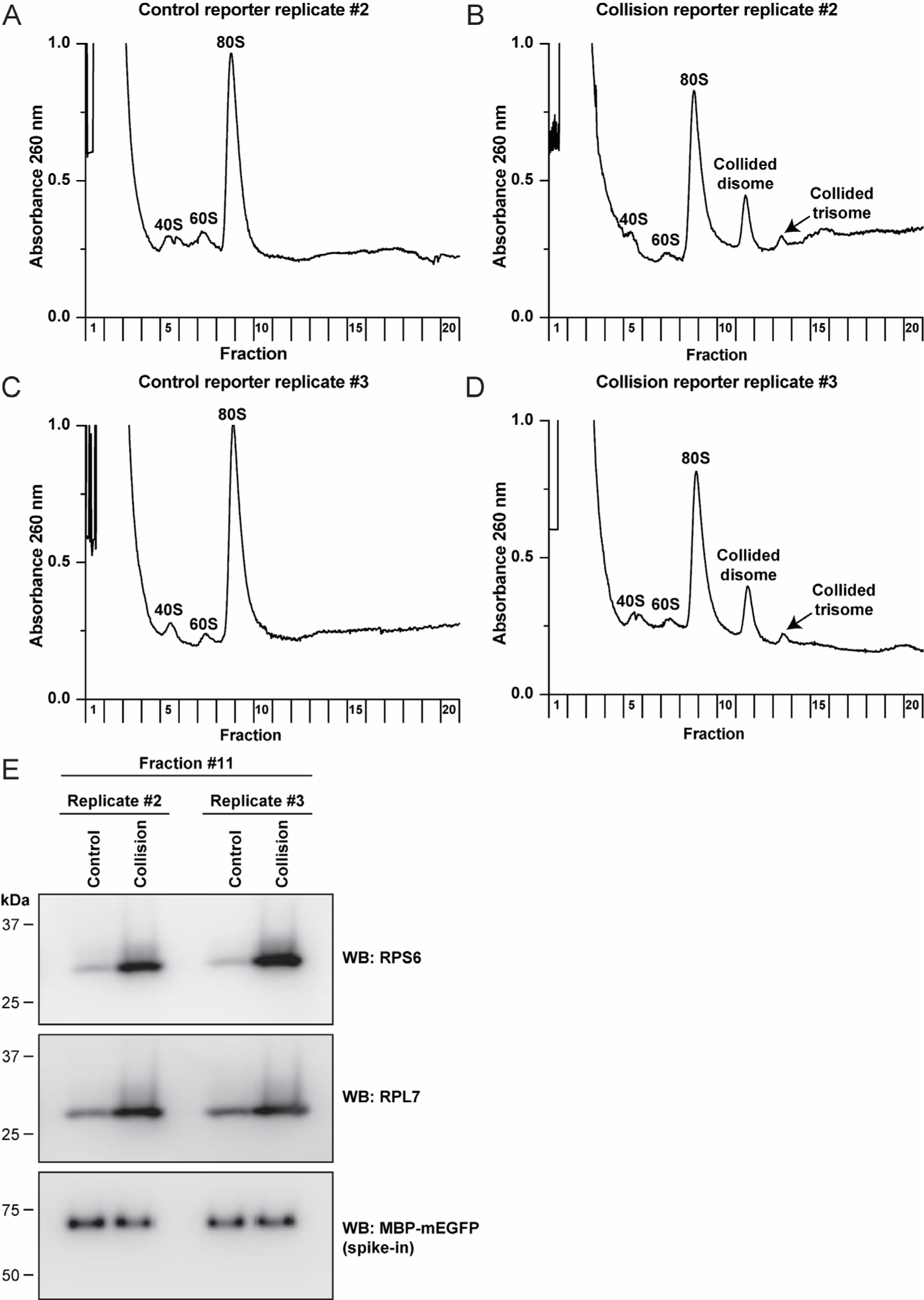
The collision reporter mRNA, but not the control reporter mRNA, generates nuclease-resistant collided ribosomes. A-D) Replicates for polysome analysis of translated control and collision reporter mRNAs with nuclease treatment related to Figure 2. The collision reporter mRNA generates nuclease-resistant collided disomes and trisomes, with a concurrent decrease in monosomes as compared to the control reporter. E) Anti-RPS6 and anti-RPL7 Western blots of fraction #11 that contains disomes. Recombinant MBP-mEGFP was spiked in and used as a loading control.

**Figure S3.**
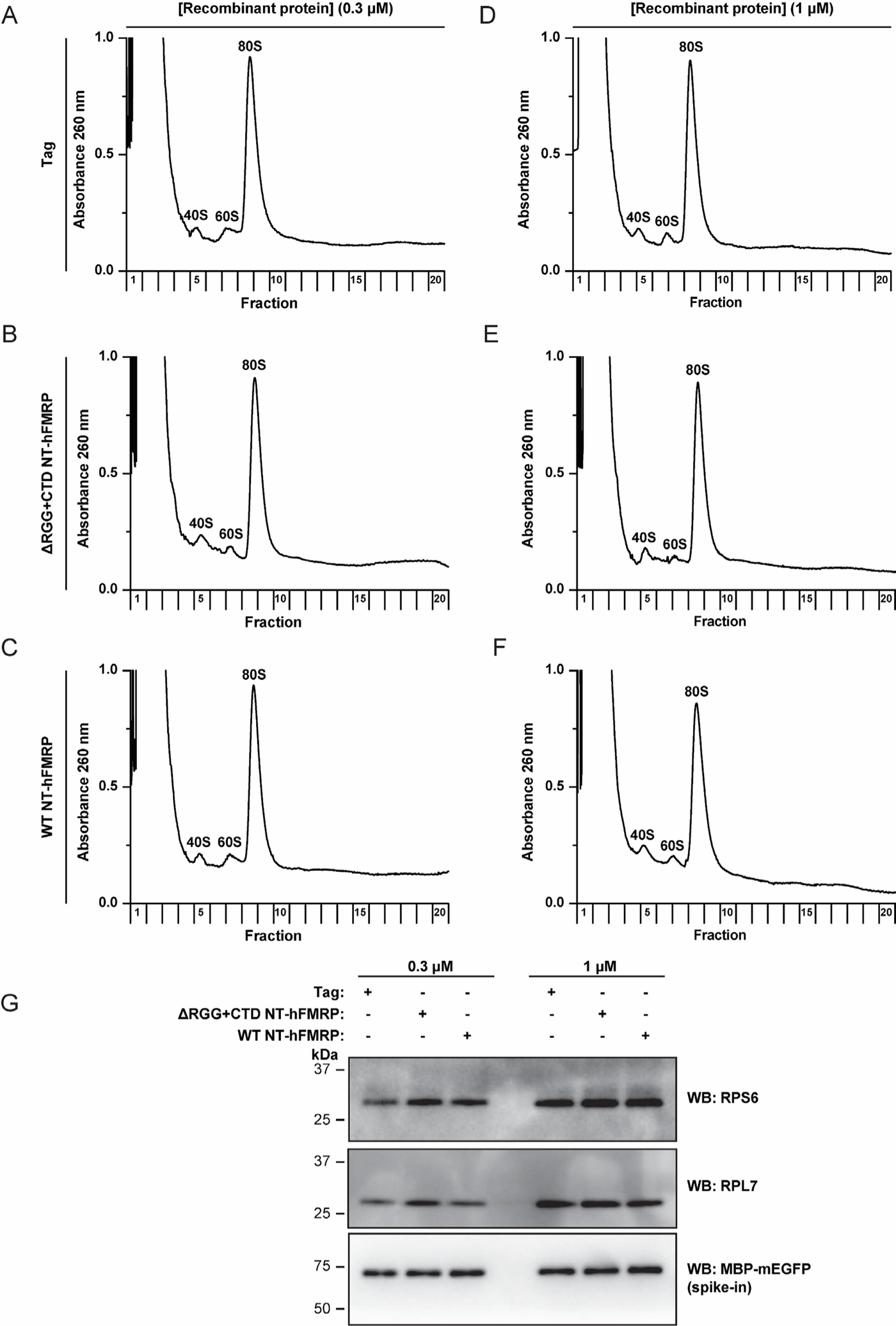
WT NT-hFMRP does not cause detectable nuclease-resistant ribosome collisions on nLuc mRNA in RRL. A-C) Polysome analysis of *in vitro* translation reactions with nuclease treatment and the indicated recombinant proteins at 0.3 μM final (A-C) or 1 µM final (D-F). G) Anti-RPS6 and anti-RPL7 Western blots of fraction #11 that contains disomes. Recombinant MBP-mEGFP was spiked in and used as a loading control.

**Figure S4.**
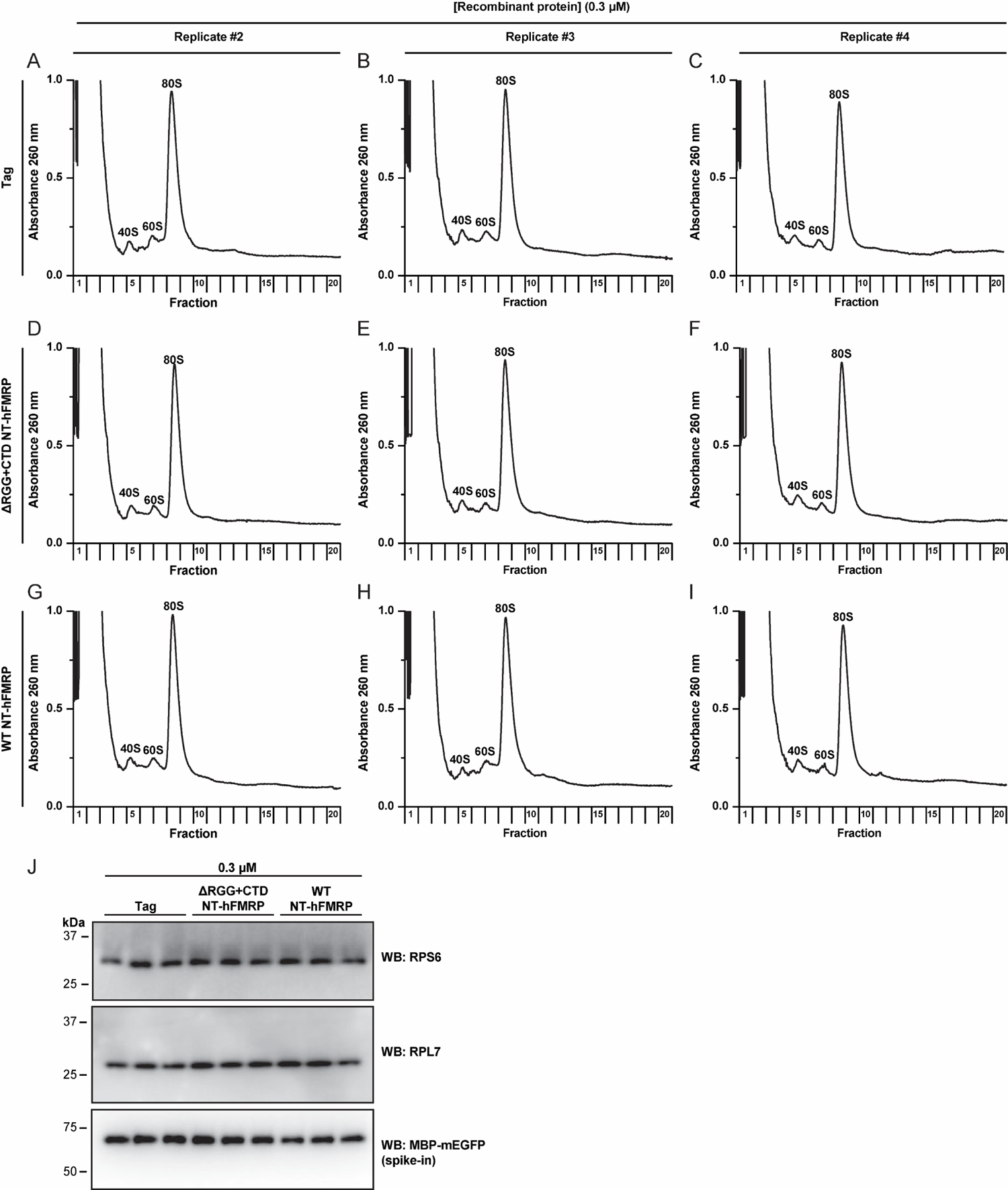
WT NT-hFMRP does not cause detectable nuclease-resistant ribosome collisions in RRL at 0.3 μM on nLuc mRNA. A-I) Replicates of polysome analysis of *in vitro* translation reactions with nuclease treatment and the indicated recombinant proteins at 0.3 μM final related to Figure S3. J) Anti-RPS6 and anti-RPL7 Western blots of fraction #11 that contains disomes. Recombinant MBP-mEGFP was spiked in and used as a loading control.

**Figure S5.**
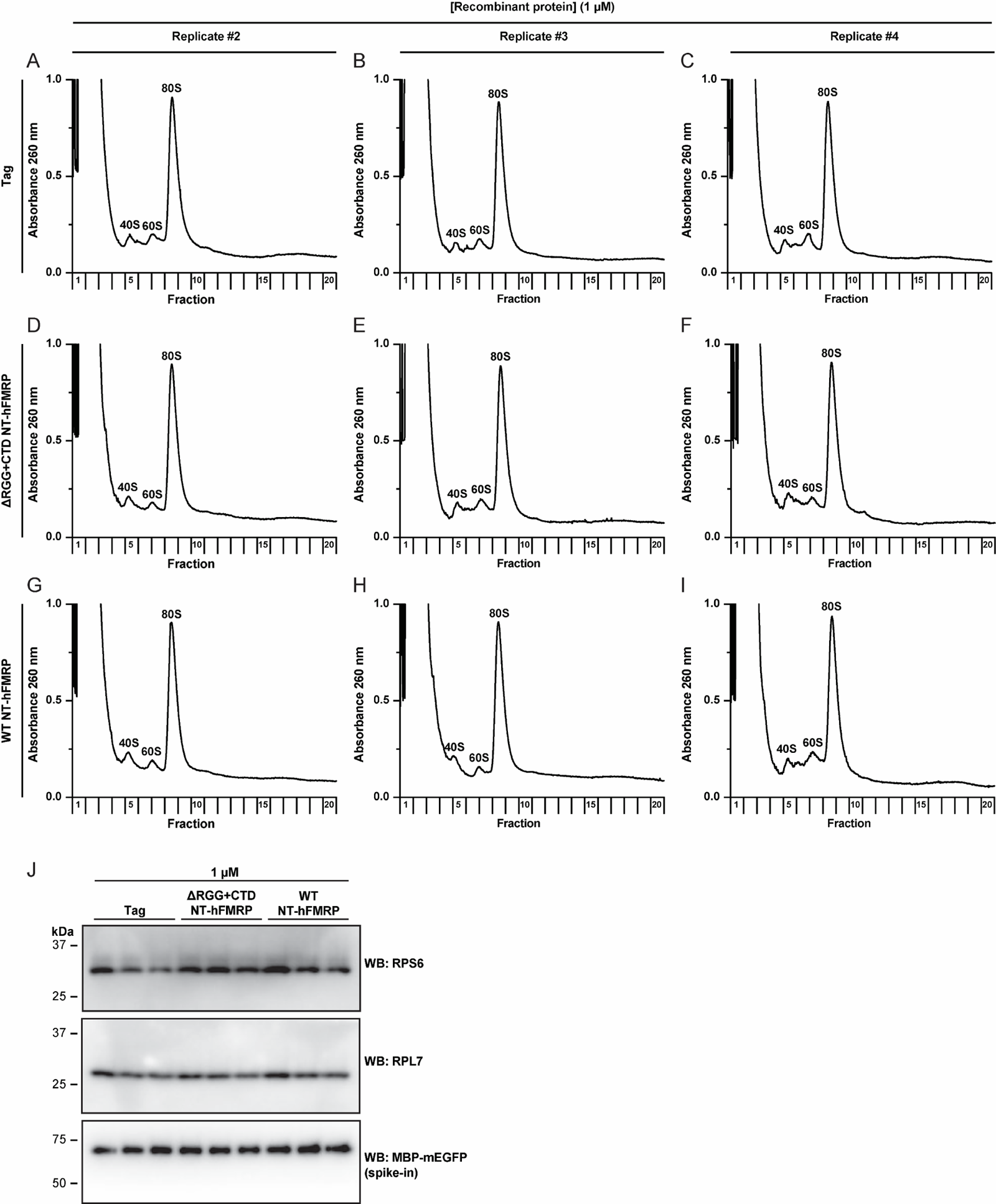
WT NT-hFMRP does not cause detectable nuclease-resistant ribosome collisions in RRL at 1 μM on nLuc mRNA. A-I) Replicates of polysome analysis of *in vitro* translation reactions with nuclease treatment and the indicated recombinant proteins at 1 μM final related to Figure S3. J) Anti-RPS6 and anti-RPL7 Western blots of fraction #11 that contains disomes. Recombinant MBP-mEGFP was spiked in and used as a loading control.

**Figure S6.**
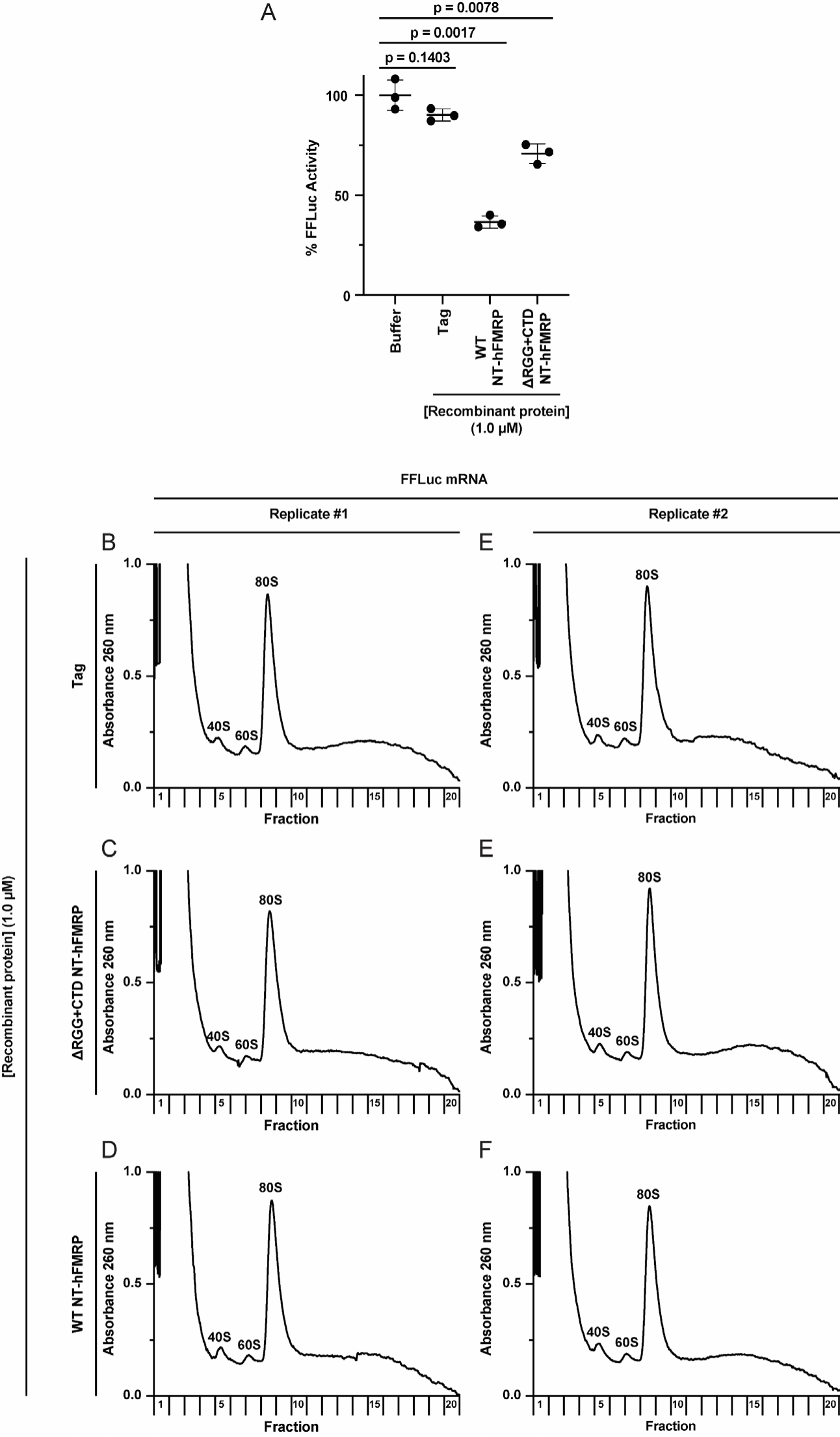
WT NT-hFMRP does not cause detectable nuclease-resistant ribosome collisions in RRL at 1 μM on FFLuc mRNA. A) *In vitro* translation of FFLuc reporter mRNAs pre-incubated with protein storage buffer (Buffer) or the indicated recombinant protein (1 µM final). Data are shown as mean ± SD. n = 3 biological replicates. Comparisons were made using a two-tailed unpaired t test with Welch’s correction. B-F) Polysome analysis of *in vitro* translation reactions with nuclease treatment and the indicated recombinant proteins at 1 μM final. Duplicate samples are shown.

**Figure S7.**
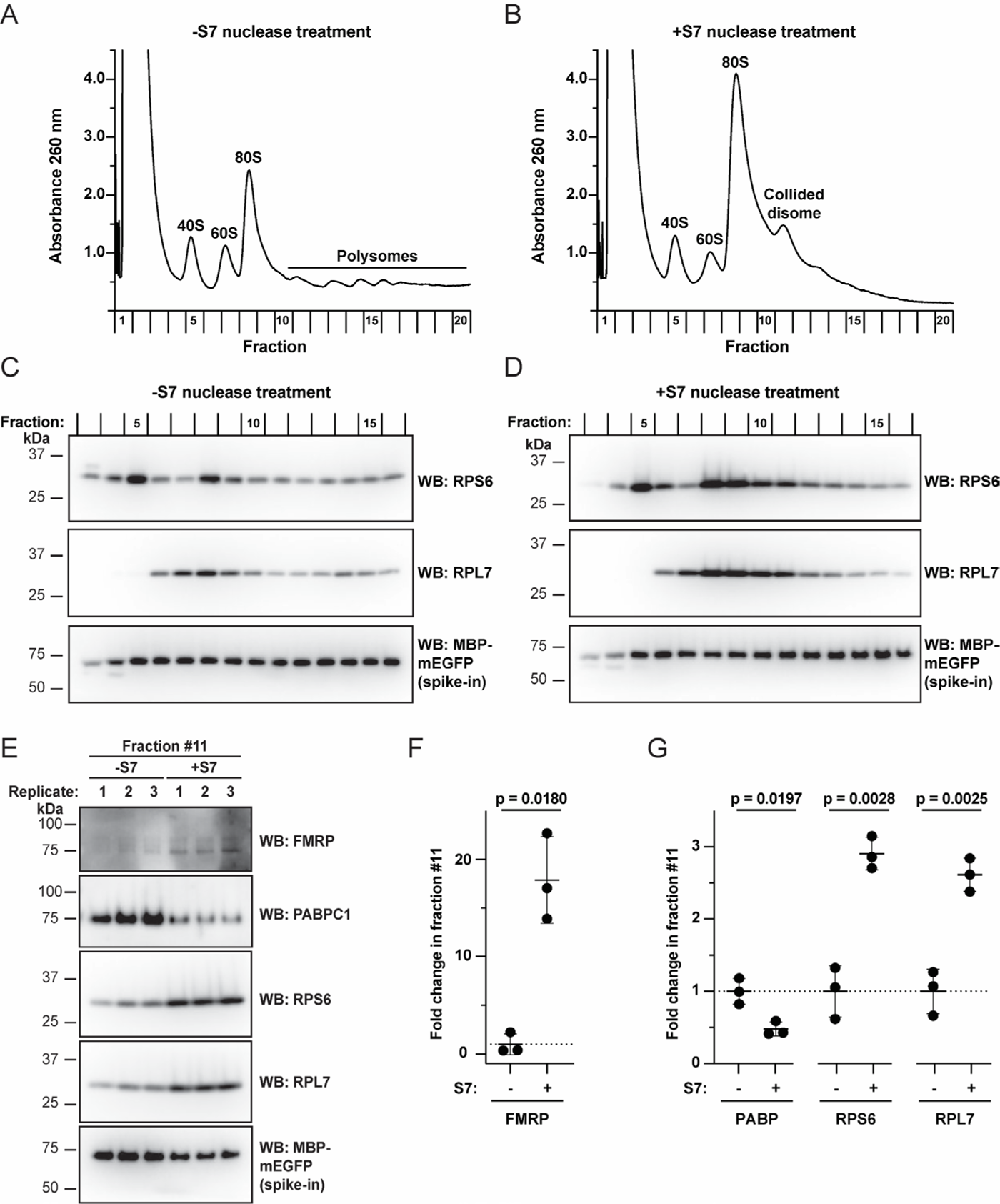
FMRP co-sediments with nuclease-resistant disomes in cells. A-B) Polysome analysis of N2A cell lysates without (A) and with (B) S7 nuclease treatment. C-D) Anti-RPS6 and anti-RPL7 Western blots of fractions 1-16 of untreated (C) or nuclease-treated cell lysates (D). The first lane contains fractions 1 & 2 combined and the second lane contains fractions 3 & 4 combined. The third lane and on contain single fractions. Recombinant MBP-mEGFP was spiked-in and used as a loading control. E) Anti-FMRP, anti-PABP, anti-RPS6, and anti-RPL7 Western blot analysis of fraction #11 that contains disomes. Recombinant MBP-mEGFP was spiked-in and used as a loading control. F-G) Quantification of the indicated proteins in panel E. Bands were first normalized to MBP-mEGFP and then set relative to untreated. Data are shown as mean ± SD. n = 3 biological replicates. Comparisons were made using a two-tailed unpaired t test with Welch’s correction.

**Figure S8.**
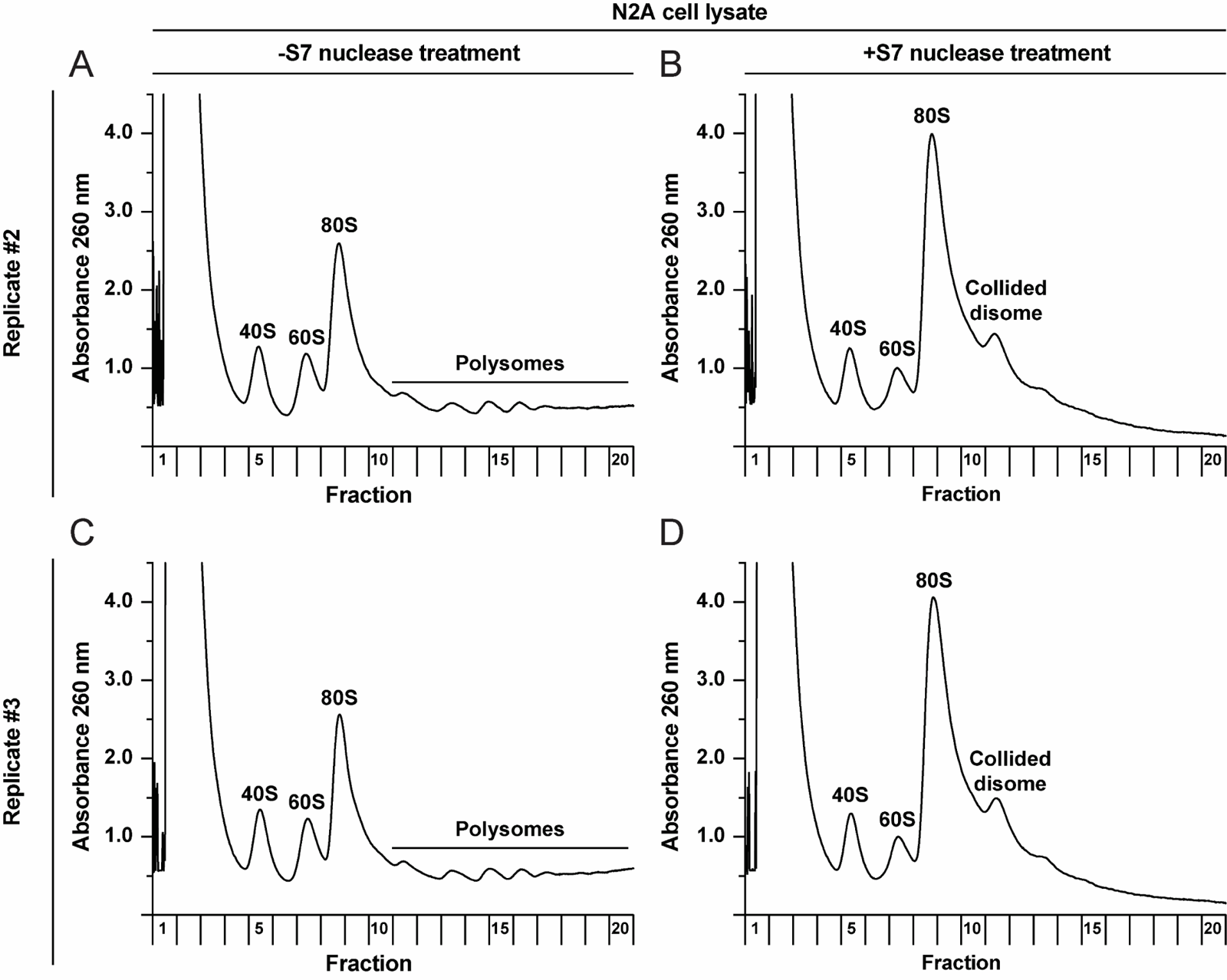
N2A cells naturally have nuclease-resistant disomes. A-D) Replicates of polysome analysis of N2A cell lysates without (A & C) and with (B & D) S7 nuclease treatment related to Figure S7.

**Figure S9.**
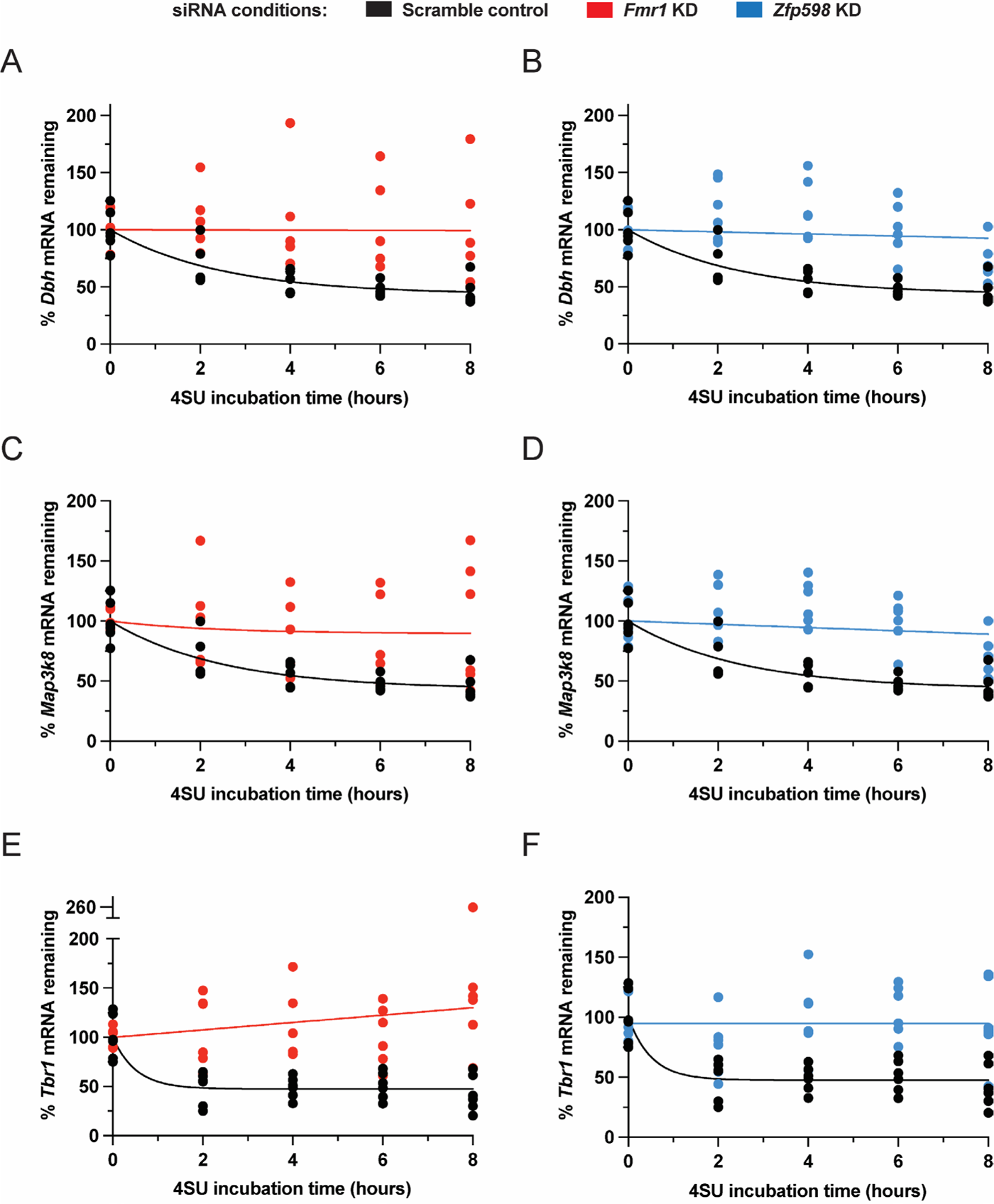
*Dbh*, *Rrh*, and *Tbr1* mRNAs are stabilized upon *Fmr1* and *Zfp598* KD compared to scramble control conditions in N2A cells. A-F) Roadblock-qPCR was used to measure mRNA half-lives (t_1/2_) of *Dbh* mRNA (A, B), *Rrh* mRNA (C, D), and *Tbr1* mRNA (E, F) in N2A cells. The Scramble negative control is shown in black (and is the same in A & B, in C & D, and in E & F), *Fmr1* KD in red, and *Zfp598* KD in blue. n=6 biological replicates. One phase decay trend lines calculated by nonlinear regression are shown. However, due to the mRNA levels not decreasing below 50% during the 8 hr time course, the t_1/2_ for these mRNAs could not be determined. Nevertheless, in all three examples, the one phase decay trend lines for both *Fmr1* KD and *Zfp598* KD are markedly above the Scramble negative contro.

---

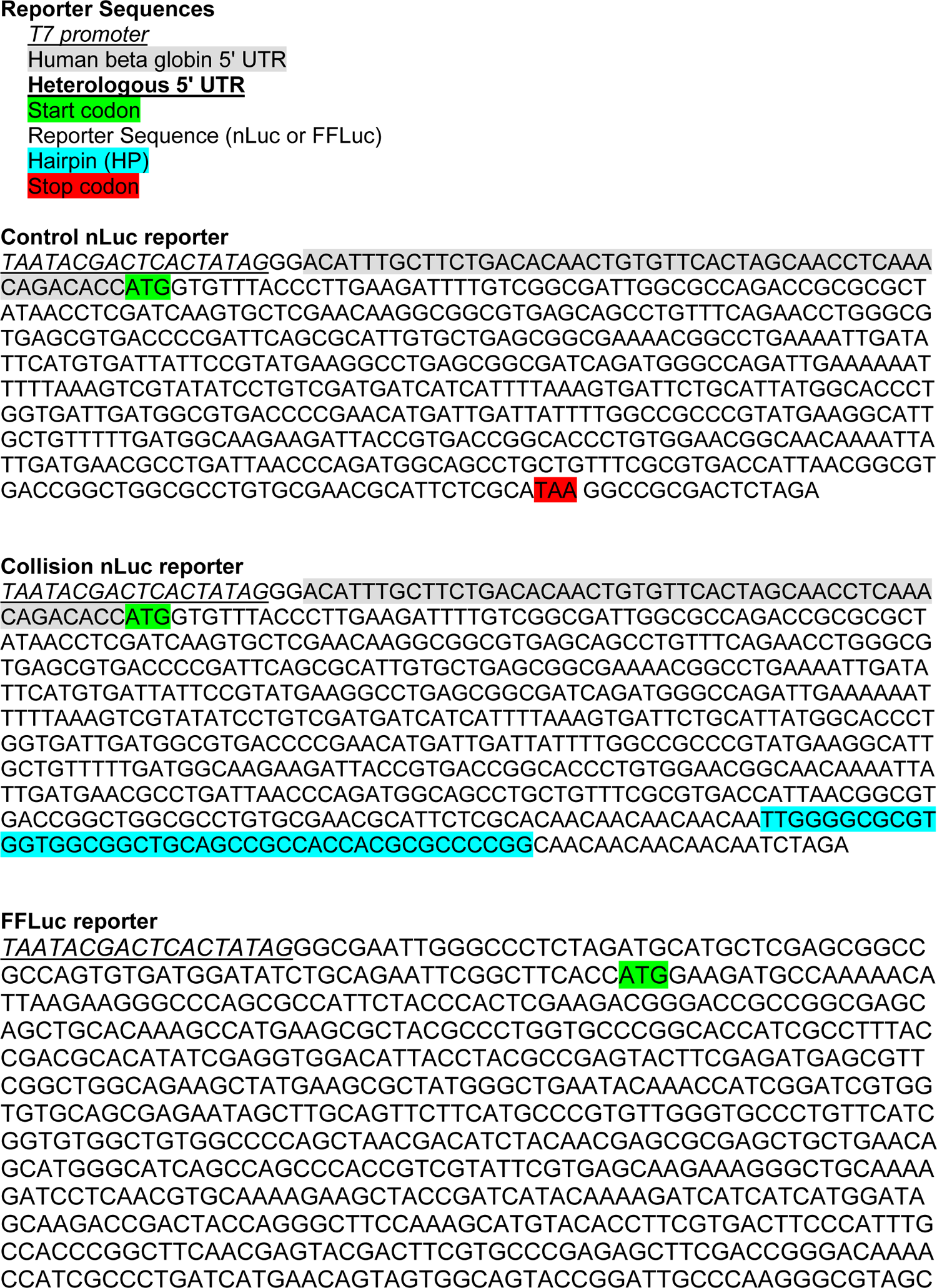
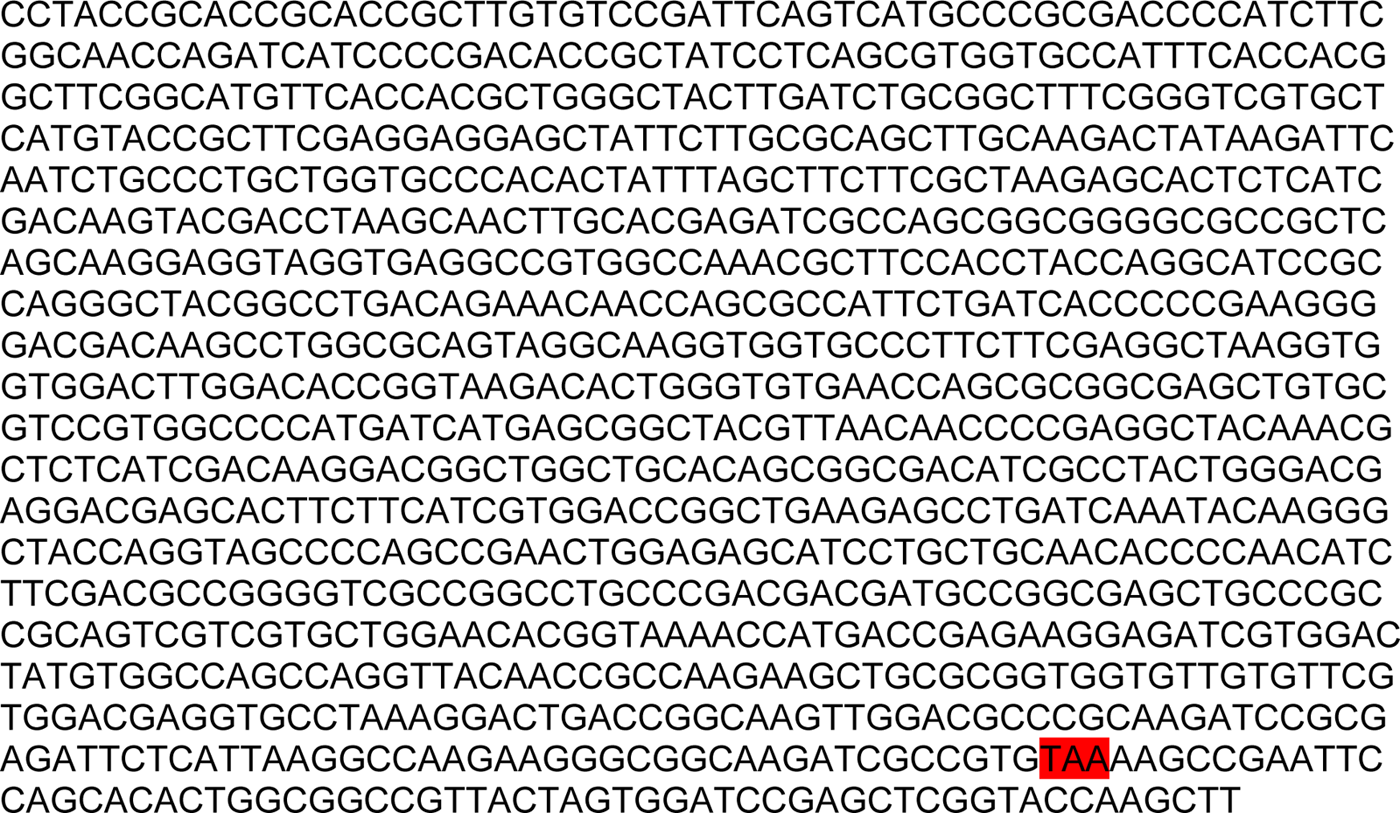
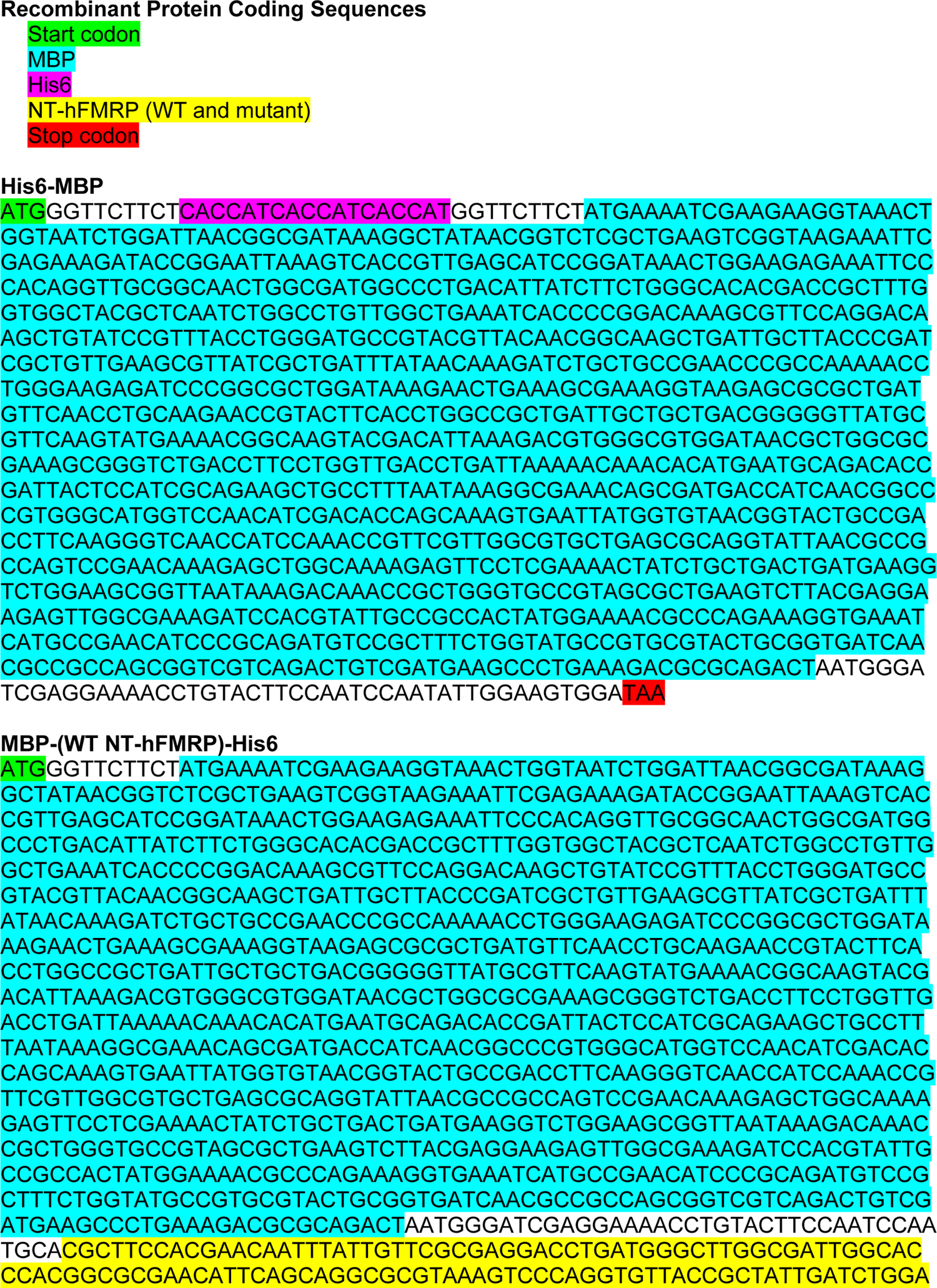
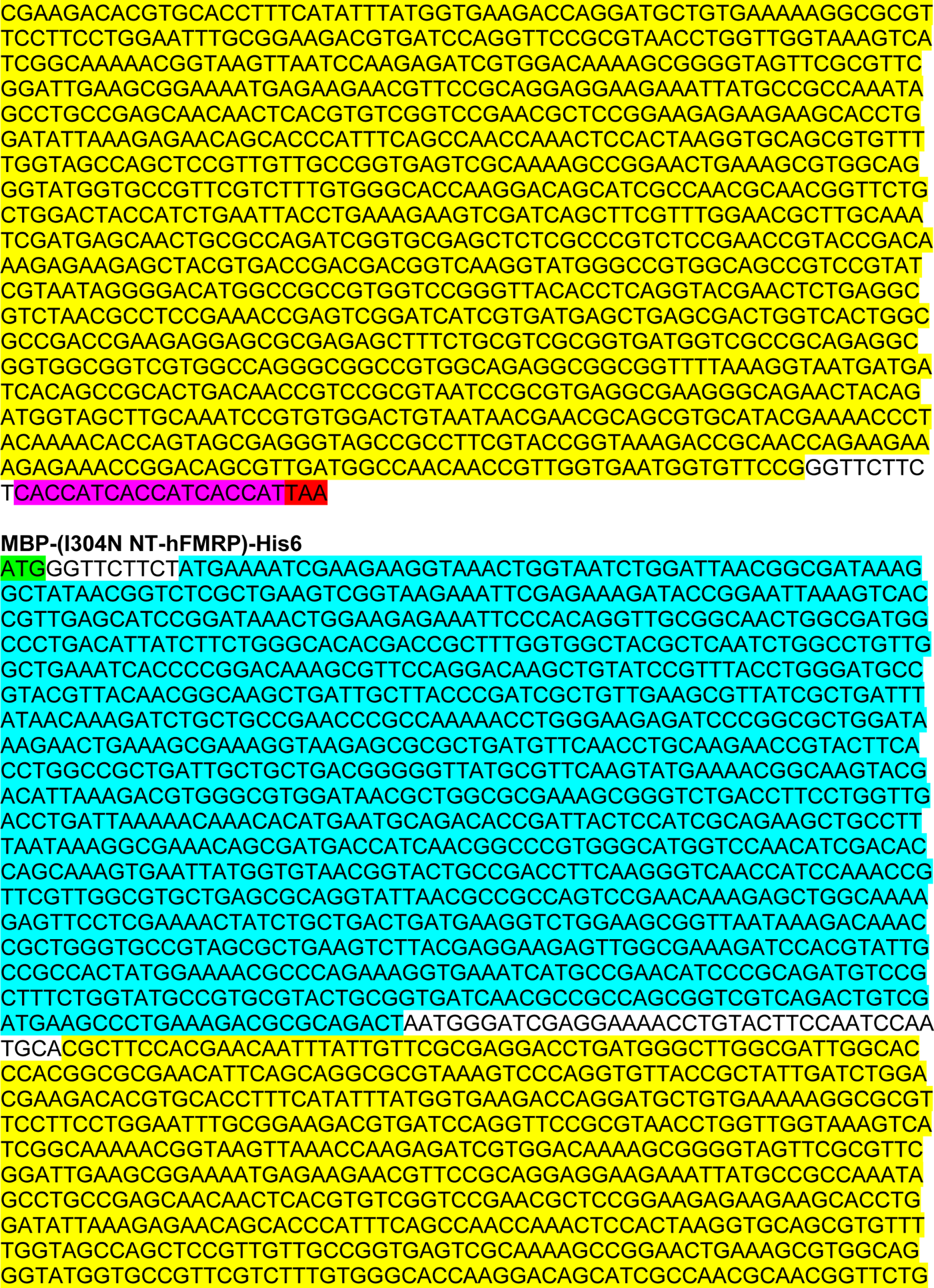
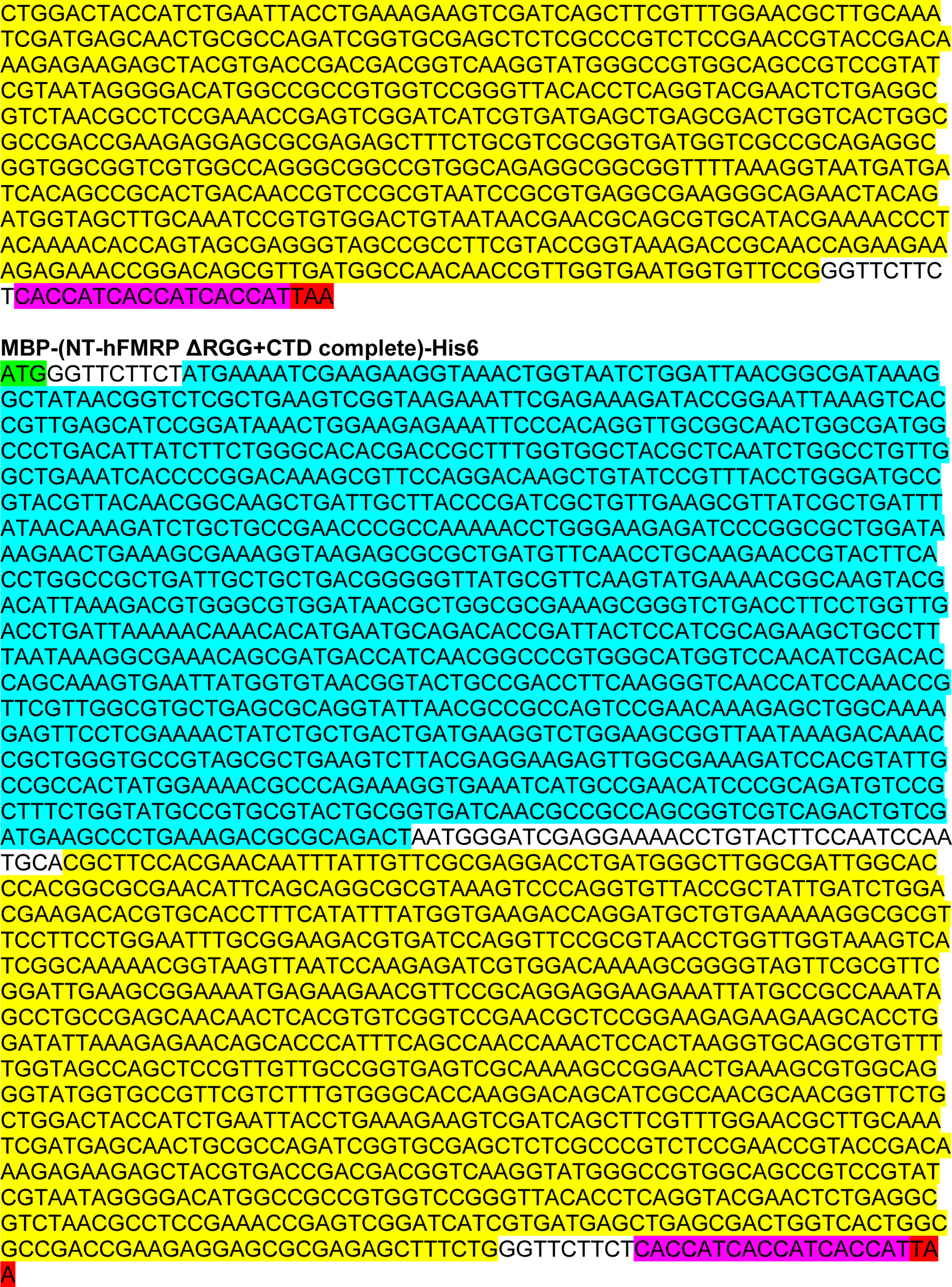
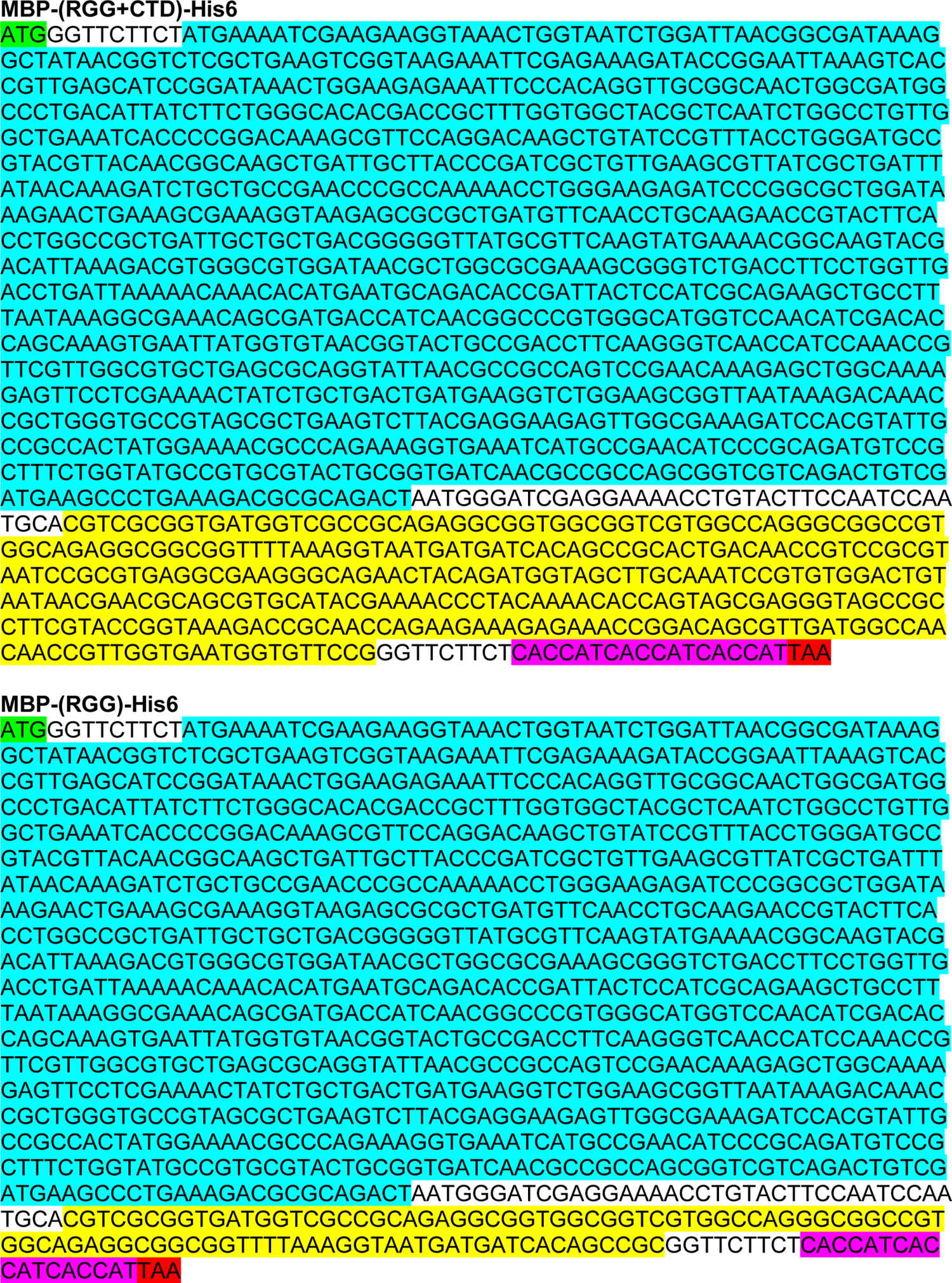
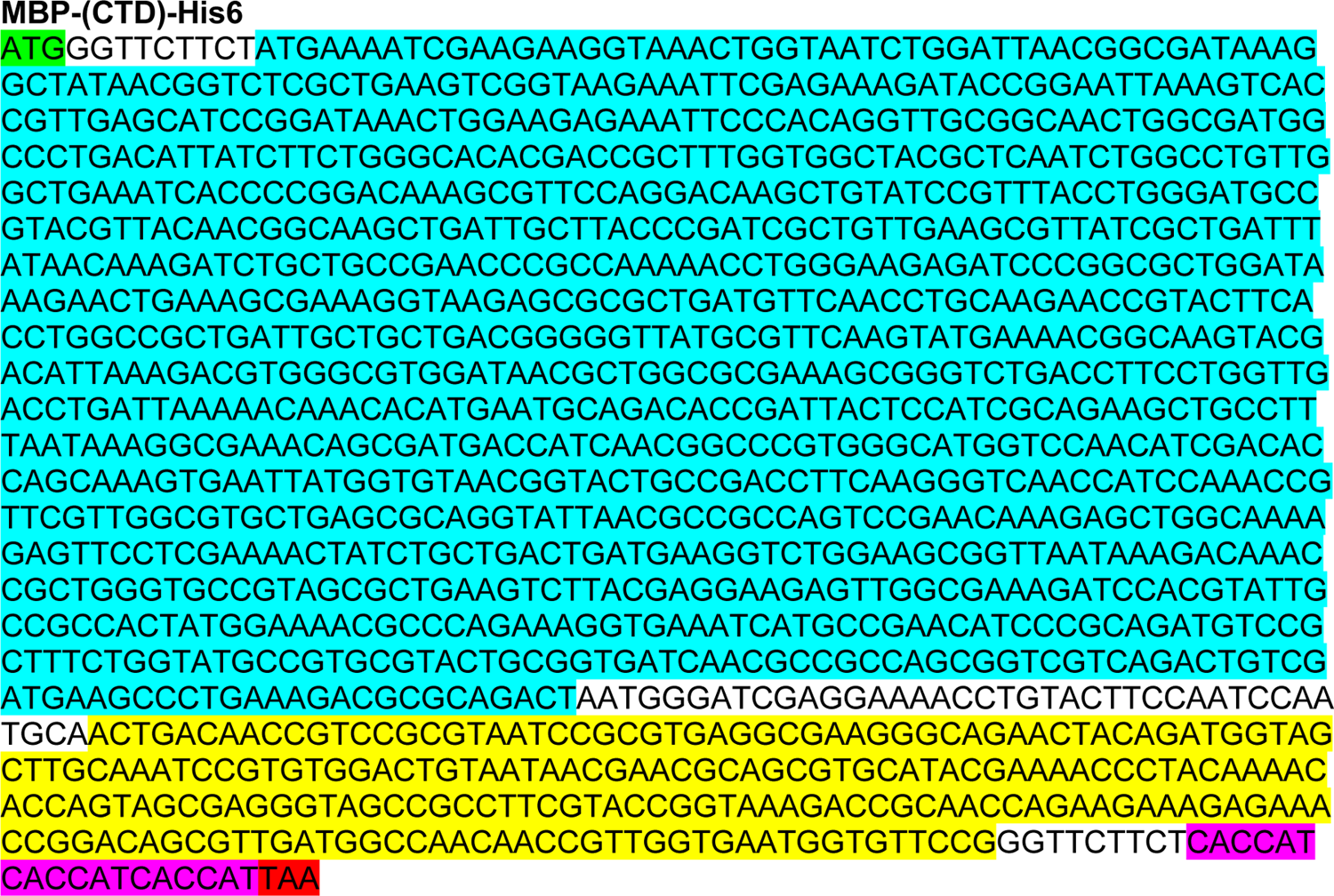

